# Keypoint-MoSeq: parsing behavior by linking point tracking to pose dynamics

**DOI:** 10.1101/2023.03.16.532307

**Authors:** Caleb Weinreb, Jonah Pearl, Sherry Lin, Mohammed Abdal Monium Osman, Libby Zhang, Sidharth Annapragada, Eli Conlin, Red Hoffman, Sofia Makowska, Winthrop F. Gillis, Maya Jay, Shaokai Ye, Alexander Mathis, Mackenzie Weygandt Mathis, Talmo Pereira, Scott W. Linderman, Sandeep Robert Datta

## Abstract

Keypoint tracking algorithms have revolutionized the analysis of animal behavior, enabling investigators to flexibly quantify behavioral dynamics from conventional video recordings obtained in a wide variety of settings. However, it remains unclear how to parse continuous keypoint data into the modules out of which behavior is organized. This challenge is particularly acute because keypoint data is susceptible to high frequency jitter that clustering algorithms can mistake for transitions between behavioral modules. Here we present keypoint-MoSeq, a machine learning-based platform for identifying behavioral modules (“syllables”) from keypoint data without human supervision. Keypoint-MoSeq uses a generative model to distinguish keypoint noise from behavior, enabling it to effectively identify syllables whose boundaries correspond to natural sub-second discontinuities inherent to mouse behavior. Keypoint-MoSeq outperforms commonly used alternative clustering methods at identifying these transitions, at capturing correlations between neural activity and behavior, and at classifying either solitary or social behaviors in accordance with human annotations. Keypoint-MoSeq therefore renders behavioral syllables and grammar accessible to the many researchers who use standard video to capture animal behavior.

## Introduction

Work from ethology demonstrates that behavior — a chain of actions traced by the body’s movement over time — is both continuous and discrete^1-3^. Keypoint tracking methods (which including SLEAP^4^, DeepLabCut^5^ and others^6,7^) enable users to specify and track points corresponding to body parts in videos of behaving animals, and thereby to quantify movement kinematics. These methods are simple to implement and applicable to a wide range of video data; because of their ease of use and generality, keypoint tracking approaches are revolutionizing our access to the continuous dynamics that underlie many aspects of animal behavior in a wide variety of settings^8^.

In contrast, it remains less clear how to best cluster behavioral data into the discrete modules of movement that serve as building blocks for more complex patterns of behavior^9-11^. Identifying these modules is essential to the creation of an ethogram, which describes the order in which behavioral modules are expressed in a particular context or experiment. While several methods exist that can automatically transform high-dimensional behavioral data into an ethogram^12-17^, their underlying logic and assumptions differ, with different methods often giving distinct descriptions of the same behavior^13,16^. An important gap therefore exists between our access to movement kinematics and our ability to understand how these kinematics are organized to impart structure upon behavior; filling this gap is essential if we are to understand how the brain builds complex patterns of action.

One widely deployed and well validated method for identifying behavioral modules and their temporal ordering is Motion Sequencing (MoSeq)^17^. MoSeq uses unsupervised machine learning methods to transform its inputs — which are not keypoints, but instead data from depth cameras that “see” in three dimensions from a single axis of view — into a set of behavioral motifs (like rears, turns and pauses) called syllables. MoSeq identifies behavioral syllables through a probabilistic generative model that instantiates the ethological hypothesis that behavior is composed of repeatedly used modules of action that are stereotyped in form and placed flexibly into at least somewhat predictable sequences. One important aspect of MoSeq is that it seeks to identify syllables by searching for discontinuities in behavioral data at a timescale that is set by the user; this timescale is specified through a “stickiness” hyperparameter that influences the frequency with which syllables can transition. In the mouse, where MoSeq has been most extensively applied, pervasive discontinuities at the sub-second-to-second timescale mark the boundaries between syllables, and the stickiness hyperparameter is explicitly set to capture this timescale. Given a timescale and a depth dataset to analyze, MoSeq automatically identifies the set of syllables out of which behavior is composed in an experiment without human supervision.

MoSeq-based analysis has captured meaningful changes in spontaneous, exploratory rodent behaviors induced by genetic mutations, changes in the sensory or physical environment, direct manipulation of neural circuits and pharmacological agents^17-20^. Importantly, MoSeq does not simply provide a useful description of behavior, but also reveals biologically important brain-behavior relationships. For example, the behavioral transitions identified by MoSeq correspond to systematic fluctuations in neural activity in both dopaminergic neurons and their targets in dorsolateral striatum (DLS)^18^, and the behavioral syllables identified by MoSeq have explicit neural correlates in DLS spiny projection neurons^19^. Furthermore, dopamine fluctuations in DLS causally influence the use and sequencing of MoSeq-identified syllables over time, and individual syllables can be reinforced (without any alteration in their underlying kinematic content) through closed-loop dopamine manipulations^18^.

However, MoSeq has a significant constraint: as currently formulated MoSeq is tailored for input data from depth cameras, which are typically placed over simple behavioral arenas in which single mice are recorded during behavior. Although depth cameras afford a high dimensional view of ongoing pose dynamics, they are also often difficult to deploy, suffer from high sensitivity to reflections, and have limited temporal resolution^21^. In principle these limits could be overcome by applying MoSeq to keypoint data. However, attempts to do so have thus far failed: researchers applying MoSeq-like models to keypoint data have reported flickering state sequences that switch much faster than the animal’s actual behavior^13,22^.

Here we confirm this finding and identify its cause: jitter in the keypoint estimates themselves, which is mistaken by MoSeq for behavioral transitions. To address this challenge, we reformulated the model underlying MoSeq to simultaneously infer correct pose dynamics (from noisy or even missing data) and the set of expressed behavioral syllables. We benchmarked the new model, called “keypoint-MoSeq”, by comparing it to both standard depth camera-based MoSeq and to alternative behavioral clustering methods (including B-SOiD^12^, VAME^13^ and MotionMapper^23^). We find that keypoint-MoSeq identifies similar sets of behavioral transitions as depth MoSeq and preserves important information about behavioral timing, despite being fed behavioral data that are relatively lower dimensional; furthermore, keypoint-MoSeq outperforms alternative methods at demarcating behavioral transitions in kinematic data, capturing systematic fluctuations in neural activity, and identifying complex features of solitary and social behavior highlighted by expert observers. Keypoint-MoSeq is flexible, and works on datasets from different labs, using overhead or bottom-up camera angles, with 2D or 3D keypoints, and in both mice and rats.

Given that keypoint tracking can be applied in diverse settings (including natural environments), requires no specialized hardware, and affords direct control over which body parts to track and at what resolution, we anticipate that keypoint-MoSeq will serve as a general tool for understanding the structure of behavior in a wide variety of settings. To facilitate broad adoption of this approach, we have built keypoint-MoSeq to be directly integrated with widely-used keypoint tracking methods (including SLEAP and DeepLabCut), and have made keypoint-MoSeq code freely accessible for academic users at www.MoSeq4all.org; this modular codebase includes novice-friendly Jupyter notebooks to enable users without extensive computational experience to use keypoint-MoSeq, methods for motif visualization in 2D and 3D, a pipeline for post-hoc analysis of the outputs of keypoint-MoSeq, and a hardware-accelerated and parallelization-enabled version of the code for analysis of large datasets.

## Results

Simple inspection of depth-based behavioral video data reveals a block-like structure organized at the sub-second timescale^17^ (Fig. 1); this observation previously inspired the development of MoSeq, which posits that these blocks encode serially-expressed behavioral syllables. To ask whether keypoint data possess a similar block-like structure, we recorded simultaneous depth and conventional two-dimensional (2D) monochrome videos at 30 Hz (using the Microsoft Azure, which has depth and IR-sensitive sensors that operate in parallel) while mice explored an open field arena; we then used a convolutional neural network to track eight keypoints in the 2D video (two ears and six points along the dorsal midline; Fig 1a).

**Figure 1:**
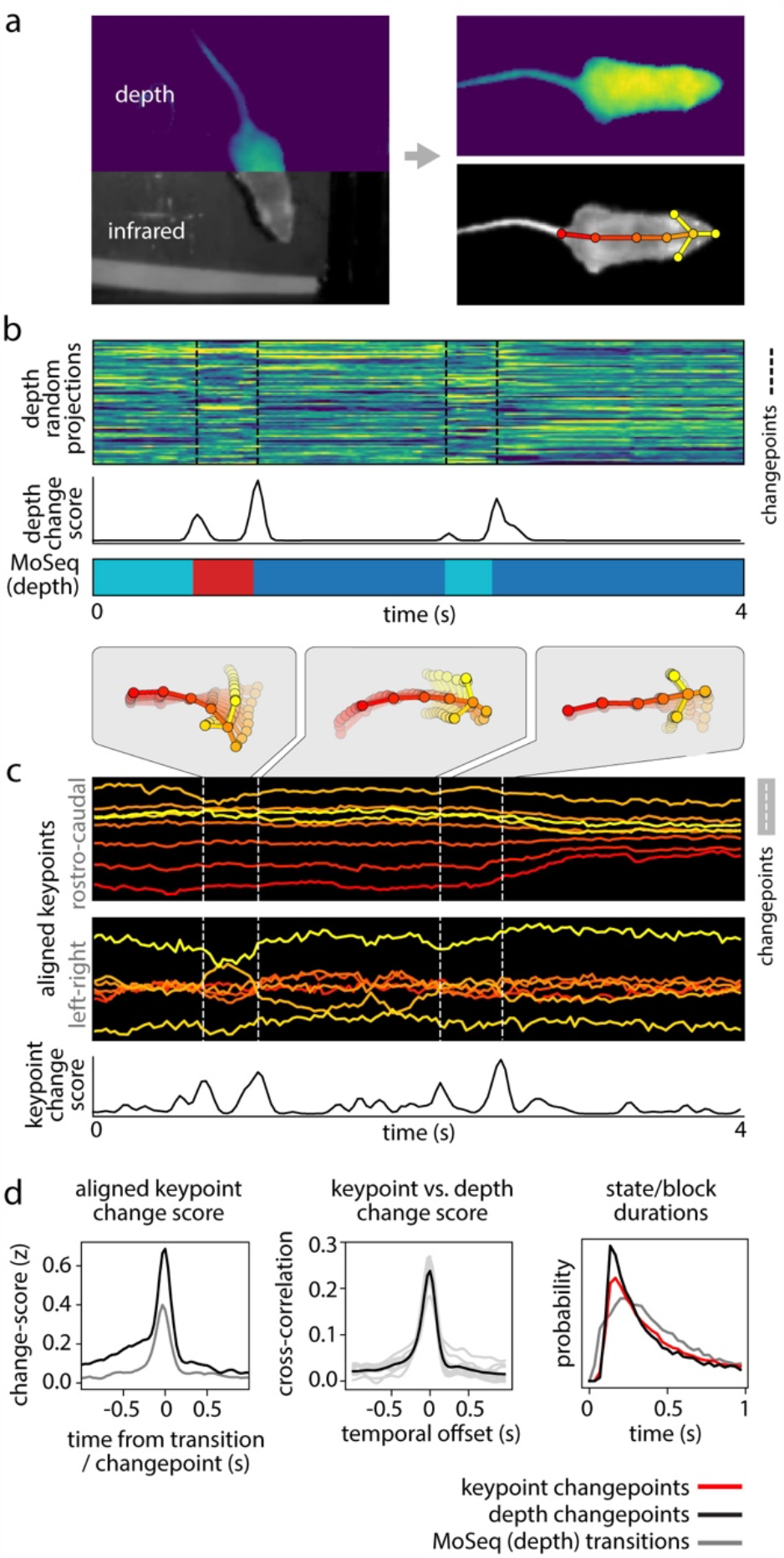
Keypoint trajectories exhibit sub-second to second structure during spontaneous behavior. **a) Left:** sample frame from simultaneous depth and 2D infrared recordings. **Right:** centered and aligned pose representations using the depth data (top) or infrared (bottom, tracked keypoints indicated). **b-c)** Features extracted from depth or 2D keypoint data within a 4-second window. All rows are temporally aligned. **b) Top:** Representation of the mouse’s pose based on depth video. Each row shows a random projection of the high-dimensional depth time-series. Discontinuities in the visual pattern capture abrupt changes in the mouse’s movement. **Middle:** Rate of change in the depth signal as quantified by a change score (see Methods). **Bottom:** color-coded syllable sequence from MoSeq applied to the depth data [referred to as “MoSeq (depth)”]. **c)** Position of each keypoint in egocentric coordinates; vertical lines mark changepoints, defined as peaks in the keypoint change score. **d) Left:** average keypoint change score (z-scored) aligned to MoSeq (depth) transitions (gray), or to changepoints in the depth signal (black). **Middle:** cross-correlation between depth- and keypoint-change scores, shown for the whole dataset (black line) and for each session (gray lines). **Right:** Distribution of syllable durations, based either on modeling or changepoint analysis.

Analysis of the depth videos (independent of MoSeq) revealed the familiar sub-second blocks of smooth behavioral dynamics punctuated by sharp transitions, and applying MoSeq to these videos segmented these blocks into a series of stereotyped behavioral syllables (Fig. 1b). Block-like structure was also apparent in the keypoint data; changepoint analysis (which identifies discontinuities in the underlying data) revealed that block durations were similar for the keypoint data, the depth data, and the syllables identified by MoSeq; furthermore, changepoints in the keypoint data matched both changepoints in the depth data and transitions in behavior identified by MoSeq (Fig 1c-d). This structure is not an accident of camera or keypoint placement, as similar results were obtained when tracking 10 keypoints (including the limbs and ventral midline) using a camera placed below the mouse (Extended Data Fig. 1). The reappearance of a common sub-second organization across depth and keypoint data suggests that this temporal structure is intrinsic to mouse behavior.

MoSeq models behavior as sequence of discrete states, where each state is defined as an autoregressive (AR) trajectory through pose space (corresponding to a syllable), and transitions between states are specified by a modified hidden Markov model (HMM). MoSeq therefore identifies syllables as repeated trajectories through pose space, and transitions between syllables as discontinuities in the pose dynamics. MoSeq includes a stickiness hyperparameter that in effect allows it to foveate on a single timescale at which it seeks to explain behavior; this feature enables MoSeq to identify syllables from depth data whose average duration is ∼400ms, although there is a broad distribution of mean durations across syllables, and each syllable is associated with its own duration distribution.

However, when applied to keypoint data, MoSeq failed to identify syllables at this characteristic ∼400ms timescale, instead producing a set of brief syllables (<100 ms) together with a small number of aberrantly long syllables that merged multiple behaviors; furthermore, the transitions between these syllables aligned poorly to changepoints derived from the keypoint data (Fig. 2a-b). These observations are consistent with prior work demonstrating that feeding keypoints to MoSeq generates behavioral representations that are less informative than those generated by alternative clustering methods^13,22^.

**Figure 2:**
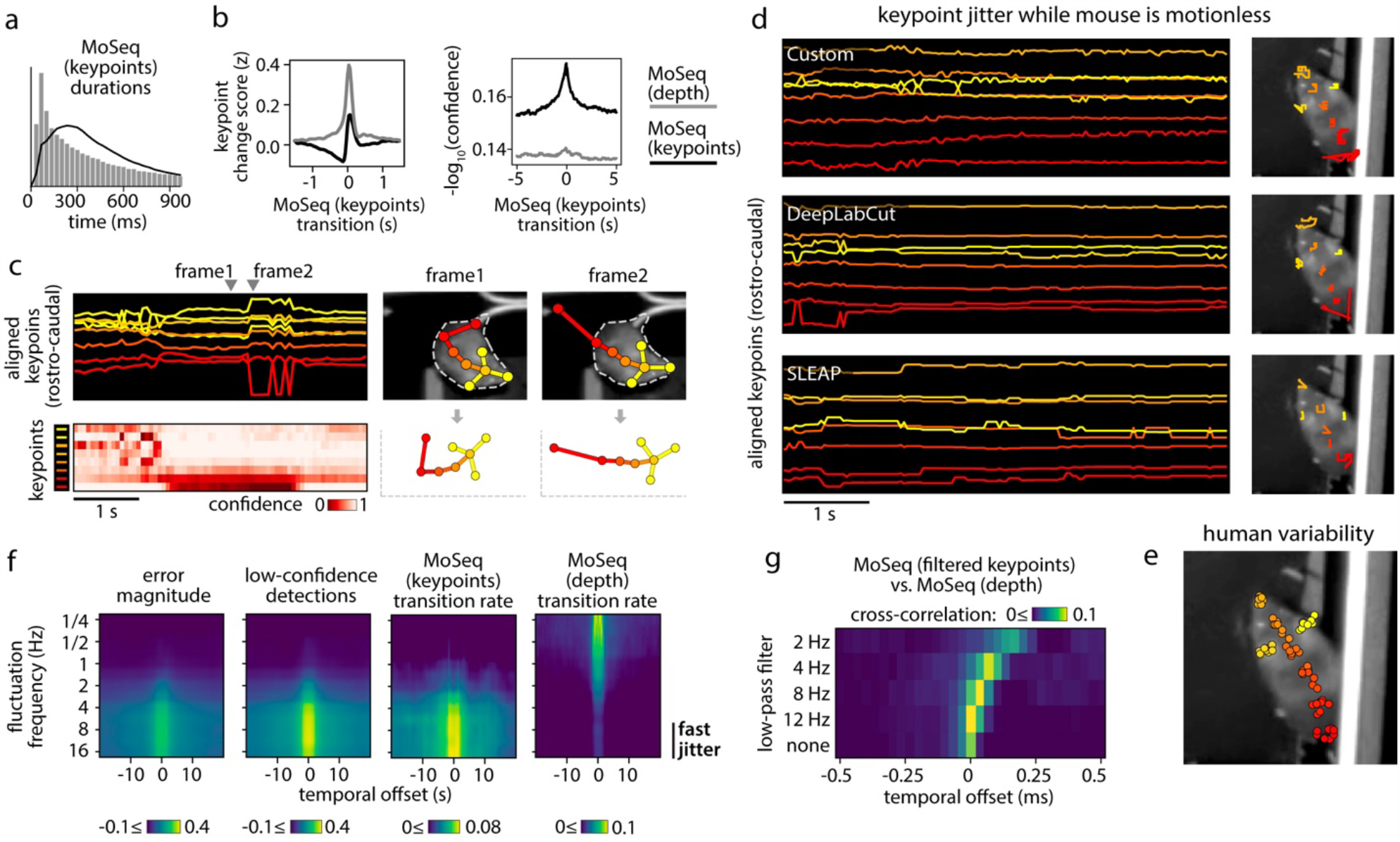
Keypoint tracking noise challenges syllable inference. **a)** Applying traditional MoSeq to keypoint trajectories [referred to as “MoSeq (keypoints)”] produces abnormally brief syllables when compared to MoSeq applied to depth data [“MoSeq (depth)”]. **b)** Keypoint change scores (left) or low-confidence detection scores (right, see Methods for how low-confidence keypoint detection was quantified), relative to the onset of MoSeq transitions (x-axis) derived from either depth (grey) or keypoint data (black). **c) Left:** example of keypoint detection errors, including high-frequency fluctuations in keypoint coordinates (top row) that coincide with low keypoint detection confidence (bottom row). **Right:** keypoint coordinates before (frame1) and during (frame2) an example keypoint position assignment error. This assignment error (occurring in the tail base keypoint) causes a shift in egocentric alignment, leading to coordinate changes across the other tracked keypoints. **d)** A five second example behavioral interval in which the same keypoints are tracked using three different methods (indicated in the inset) reveal pervasive jitter during stillness. **Left:** egocentrically aligned keypoint trajectories. **Right:** path traced by each keypoint during the 5-second interval. **e)** Variability in keypoint positions assigned by eight human labelers (see Methods). **f)** Cross-correlation between various features and keypoint fluctuations at a range of frequencies. Each heatmap represents a different scalar time-series (such as “transition rate” – the likelihood of a syllable transition on each frame), each row shows the cross-correlation between that time-series and the time-varying power of keypoint fluctuations at a given frequency. **g)** Timing of syllable transitions when MoSeq is applied to smoothed keypoint data, from most smoothed (top) to least smoothed (bottom). Each row shows the cross-correlation of MoSeq transition rates between keypoints and depth (i.e., the relative timing and degree of overlap between syllable transitions from each model).

We wondered whether the poor performance of MoSeq could be explained by noise in the keypoint data, which in principle could introduce subtle discontinuities that are falsely recognized by MoSeq as behavioral transitions. Indeed, mouse keypoint data exhibited high-frequency (>8Hz) jitter in position regardless of whether we tracked keypoints with our custom neural network or with commonly used platforms like DeepLabCut (DLC) and SLEAP (Fig. 2c-d, see Methods). Inspection of videos revealed that high frequency keypoint jitter is often associated with local tracking errors or rapid switching in the inferred location of an ambiguously positioned keypoint, rather than discernable changes in pose (Fig 2d, Extended Data Fig. 2a, Suppl. Movie 1). Indeed, frame-to-frame fluctuations in the keypoints had a similar scale as the variability in human labeling and as the test error in heldout image annotations (Fig 2e, Extended Data Fig. 2b-d). We confirmed that keypoint flicker was unrelated to true movement by tracking the same body part using multiple cameras; though overall movement trajectories were almost identical across cameras, the high-frequency fluctuations around those trajectories were uncorrelated, suggesting that the fluctuations are an artifact of tracking (Extended Data Fig. 2e-f). Consistent with the possibility that keypoint noise dominates MoSeq’s view of behavior, syllable transitions derived from keypoints – but not depth – frequently overlapped with jitter and low-confidence estimates of keypoint position (Fig. 2f). Though one might imagine that simple smoothing could ameliorate this problem, application of a low-pass filter had the additional consequence of blurring actual transitions, preventing MoSeq from identifying syllable boundaries (Fig 2g). Median filtering and Gaussian smoothing similarly yielded no improvement (Extended Data Fig 2g). These data reveal that high-frequency tracking noise can be pervasive across point-tracking algorithms and demonstrate that this noise impedes the ability of MoSeq to accurately segment behavior.

### Hierarchical modeling of keypoint trajectories decouples noise from behavior

MoSeq syllables reflect keypoint jitter because MoSeq assumes that each keypoint is a faithful and accurate representation of the position of a point on the animal. We therefore sought an alternative approach that could treat the keypoints as noisy observations rather than the truth. Switching linear dynamical systems (SLDS), which extend the AR-HMM model that underlies MoSeq, offer a principled way to decouple keypoint noise from behavior^24,25^. We therefore formulated an SLDS-based version of MoSeq whose architecture enables joint inference of pose and syllable structure. This new SLDS model has three hierarchical levels (Fig. 3a): a discrete state sequence (top level) that governs the dynamics of keypoint trajectories in a low-dimensional pose space (middle level), which is then projected into the keypoint space itself (bottom level). The three levels of this model therefore correspond to syllables, pose dynamics, and keypoint coordinates respectively.

**Figure 3:**
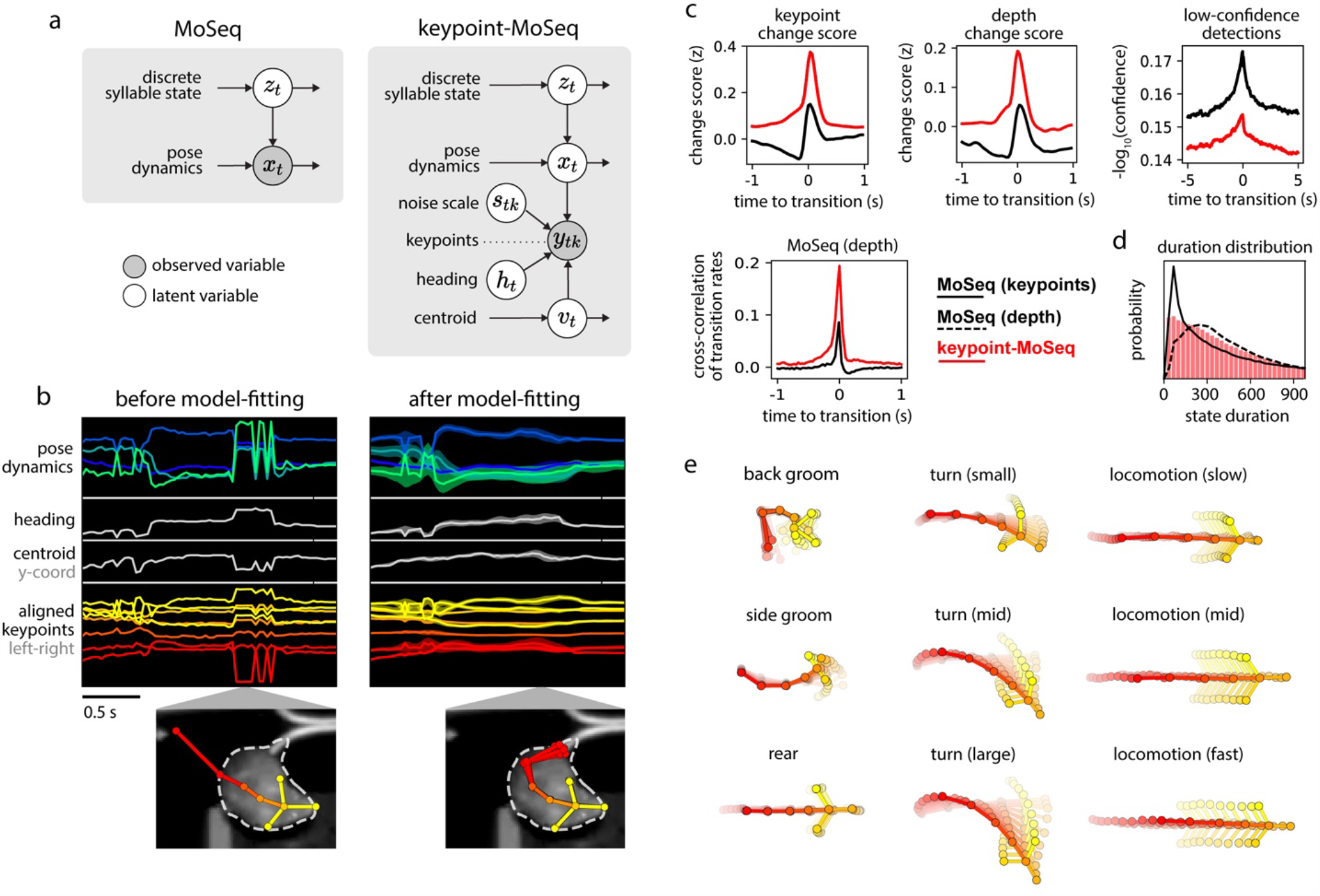
Hierarchical modeling of keypoint trajectories decouples noise from pose dynamics. **a)** Graphical models illustrating traditional and keypoint-MoSeq. In both models, a discrete syllable sequence governs pose dynamics; these pose dynamics are either described using PCA (as in “MoSeq”, left) or are inferred from keypoint observations in conjunction with the animal’s centroid and heading, as well as a noise scale that discounts keypoint detection errors (as in “keypoint-MoSeq”, right). **b)** Example of error correction by keypoint-MoSeq. **Left:** Before fitting, all variables (y axis) are perturbed by incorrect positional assignment of the tail base keypoint (whose erroneous location is shown in the bottom inset). **Right:** Keypoint-MoSeq infers plausible trajectories for each variable (shading represents the 95% confidence interval). The inset shows several likely keypoint coordinates for the tail base inferred by the model. **c) Top:** Average values of various features aligned to syllable transitions from keypoint-MoSeq (red) vs. traditional MoSeq applied to keypoint data (black). Bottom: cross-correlation of syllable transition rates between each model and depth MoSeq. Peak height represents the relative frequency of overlap in syllable transitions. **d)** Duration distribution of the syllables from each of the indicated models. **e)** Average pose trajectories for example keypoint-MoSeq syllables. Each trajectory includes ten poses, starting 165ms before and ending 500ms after syllable onset.

We further adapted the SLDS model to keypoint data by adding three additional variables: centroid and heading (which capture the animal’s overall position in allocentric coordinates) and a noise estimate for each keypoint in each frame^26^. When fit to data, the SLDS model estimates for each frame the animal’s location and pose, as well as the identity and content of the current behavioral syllable (Fig. 3b). Because of its structure, when a single keypoint implausibly jumps from one location to another, the SLDS model can attribute the sudden displacement to noise and preserve a smooth pose trajectory; if all the keypoints suddenly rotate within the egocentric reference frame, the model can adjust the inferred heading for that frame and restore a plausible sequence of coordinates. Since in the special case of zero keypoint noise our new model reduces to the same AR-HMM used in depth MoSeq^17^, we refer to this new method as “keypoint-MoSeq” for the remainder of the paper.

Unlike traditional MoSeq, keypoint-MoSeq appeared to effectively identify behavioral syllables rather than noise in the keypoint data. State transitions identified by keypoint-MoSeq overlapped with transitions in the raw depth data, with depth MoSeq-derived syllable transitions, and with transitions in the keypoints as identified by changepoint analysis; syllable boundaries identified by keypoint-MoSeq also overlapped less with low-confidence neural network detections for individual keypoints (Fig. 3c). Furthermore, the duration distribution of syllables identified by keypoint-MoSeq more closely matched that generated by conventional MoSeq using depth data (Fig 3d, Extended Data Fig 3a). From a modeling perspective the output of MoSeq was sensible: cross-likelihood analysis revealed that keypoint-based syllables were mathematically distinct trajectories in pose space, and submitting synthetic keypoint data that lacked any underlying block structure resulted in keypoint-MoSeq models that failed to identify distinct syllables (Extended Data Fig 3b,c). These analyses suggest that keypoint-MoSeq effectively addresses the syllable switching problem, nominating it as a candidate for parsing keypoint data obtained from conventional 2D cameras into syllables.

For our open field data, keypoint-MoSeq identified 25 syllables (Extended Data Fig 3d, Suppl Movie 2). Inspection of movies depicting multiple instances of the same syllable revealed that each syllable was a distinct, stereotyped motif of behavior that could be easily labeled by human observers (Suppl Movie 3). Keypoint-MoSeq differentiated between categories of behavior (e.g., rearing, grooming, walking), and variations within each category (e.g., turn angle, speed) (Fig 3e). Importantly, keypoint-MoSeq preserves access to the kinematic and morphological parameters that underlie each behavioral syllable (Extended Data Fig 3e), thereby enabling explicit comparisons and analysis. These data demonstrate that keypoint-MoSeq provides an interpretable segmentation of behavior captured by standard 2D videos, which are used in most behavioral neuroscience experiments.

### Keypoint-MoSeq better captures the fast temporal structure of behavior than alternative behavioral clustering methods

Although there is no single agreed-upon metric that can be used to validate an unsupervised segmentation of behavior, we reasoned that keypoint-MoSeq would be useful to behavioral neuroscientists if it identified boundaries between behavioral states that correspond to recognizable transitions in animal behavior, and if its outputs meaningfully relate to neural activity. As part of the validation process we also compared keypoint-MoSeq to alternative unsupervised methods for clustering keypoints, in the hopes that these comparisons might highlight strengths and weaknesses that are particular to each method. Such alternative methods include VAME, MotionMapper and B-SOiD, all of which first transform keypoint data into a feature space that reflects the dynamics in a small window around each frame, and then cluster those features to distinguish a set of behavioral states^12,13,23,27^.

As mentioned above, by design MoSeq identifies boundaries between behavioral syllables that correspond to abrupt transitions in the keypoint or depth data. To ask whether alternative behavioral clustering methods identify similar boundaries between discrete behaviors, we applied them to the identical 2D keypoint dataset. Behavioral states from VAME, B-SOiD and MotionMapper were usually brief (median duration 33-100ms, compared to ∼400ms for keypoint-MoSeq) and their transitions aligned significantly less closely with changepoints in keypoint data than did syllable transitions identified by keypoint-MoSeq (Fig 4a-c). To ensure these results were the consequence of the methods themselves rather than specific parameters we chose, we performed a comprehensive parameter scan for all methods, including up to an order of magnitude dilation of the temporal windows used by B-SOiD and MotionMapper, as well as scans over latent dimension, state number, clustering mode, and preprocessing options across all methods (where applicable); this analysis revealed some parameter combinations that yielded longer state durations, but these combinations tended to have a similar or worse alignment to changepoints in the keypoint data, a finding we replicated for both overhead and bottom-up camera angles (Extended Data Figure 4a).

**Figure 4:**
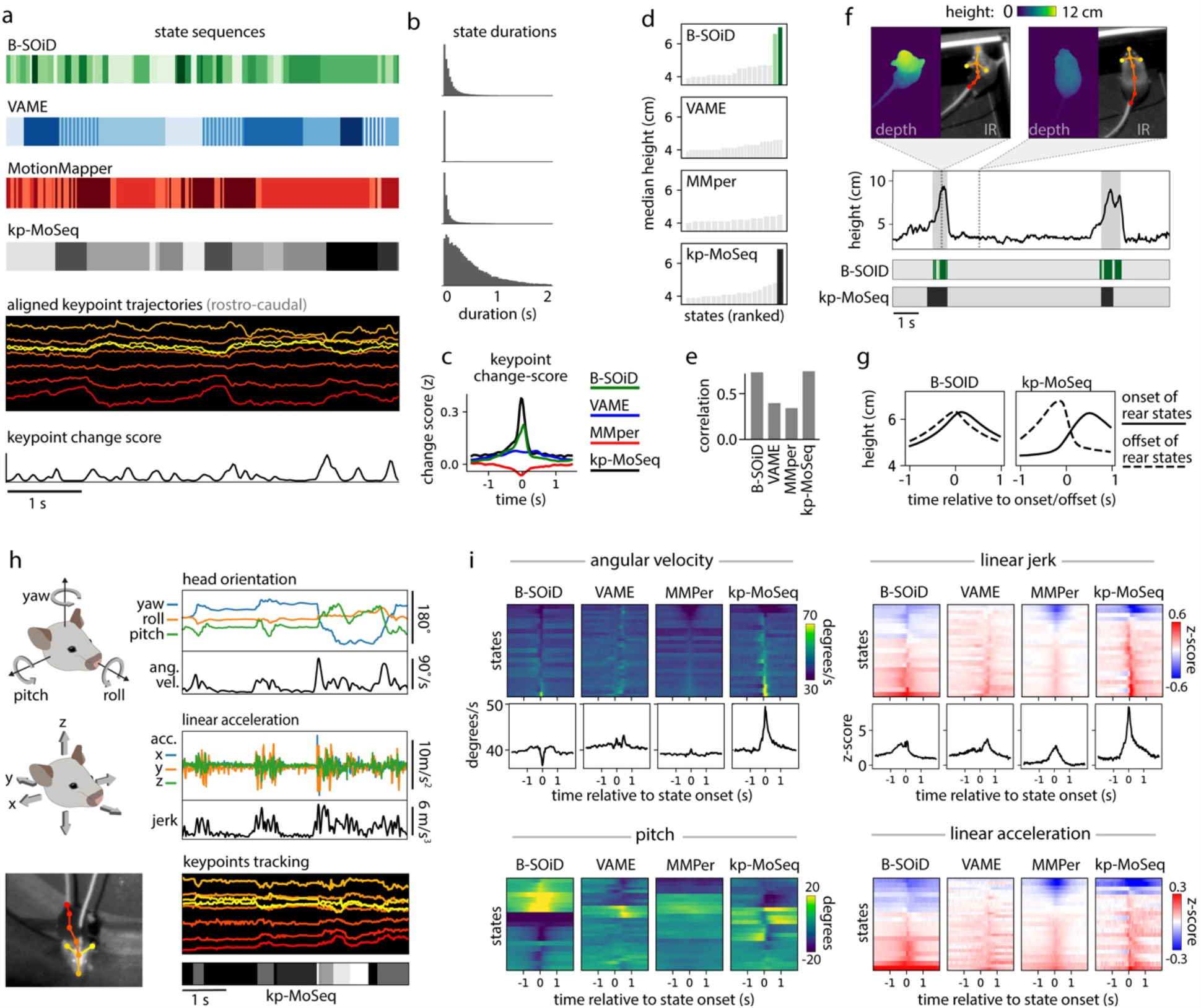
Keypoint-MoSeq captures the temporal structure of behavior. **a)** Example behavioral segmentations from four methods applied to the same 2D keypoint dataset. Keypoint-MoSeq transitions (fourth row) are sparser than those from other methods and more closely aligned to peaks in keypoint change scores (bottom row). **b)** Distribution of state durations for each method in (a). **c)** Average keypoint change scores (z-scored) relative to transitions identified by the indicated method (“MMper” refers to MotionMapper). **d)** Median mouse height (measured by depth camera) for each unsupervised behavior state. Rear-specific states (shaded bars) are defined as those with median height > 6cm. **e)** Accuracy of models designed to decode mouse height, each of which were fit to state sequences from each of the indicated methods. **f) Bottom:** state sequences from keypoint-MoSeq and B-SOiD during a pair of example rears. States are colored as in (d). **Top:** mouse height over time with rears shaded gray. Callouts show depth- and IR-views of the mouse during two example frames. **g)** Average mouse height aligned to the onsets (solid line) or offsets (dashed line) of rear-specific states defined in (d). **h)** Signals captured from a head-mounted inertial measurement unit (IMU), including absolute 3D head-orientation (top) and relative linear acceleration (bottom). Each signal and its rate of change, including angular velocity (ang. vel.) and jerk (the derivative of acceleration), is plotted during a five second interval. **i)** IMU signals aligned to the onsets of each behavioral state. Each heatmap row represents a state. Line plots show the median across states for angular velocity and jerk.

Rearing affords a particularly clear example of the differences between unsupervised behavioral methods with respect to time. B-SOiD and keypoint-MoSeq both learned a specific set of rear states/syllables from 2D keypoint data (Fig 4d; no rear-specific states were identified by VAME or MotionMapper) and each encoded the mouse’s height with comparable accuracy (B-SOiD: R=0.73, keypoint-MoSeq: R=0.74 for correlation between predicted and true mouse height; Fig 4e). Yet the rear states from each method differed dramatically in their dynamics. Whereas keypoint-MoSeq typically detected two syllable transitions that surrounded each rearing behavior (one entering the rearing syllable, the second exiting the rearing syllable), B-SOiD typically detected five to ten different transitions during the execution of a single rear, including switches between distinct rear states as well as flickering between rear- and non-rear-states (Fig 4f; Extended Data Fig 4b). This difference was made further apparent when we aligned mouse height to rearing states identified by the different methods (Fig 4g). Mouse height increased at transitions into keypoint-MoSeq’s rear state and fell at transitions out of it, producing a pair of height trajectories into and out of the rearing syllable that differed from each other and were asymmetric in time. In contrast, height tended to peak at transitions into and out of B-SOiD’s rear states, with a temporally symmetric trajectory that was only slightly different for ingoing versus outgoing transitions; this observation suggests that — at least in this example — B-SOiD does not effectively identify the boundaries between syllables, but instead tends to fragment sub-second behaviors throughout their execution.

The observation that keypoint-MoSeq effectively identifies behavioral boundaries has so far relied exclusively on analysis of video data. We therefore sought to validate keypoint-MoSeq and compare it to other unsupervised behavioral algorithms using a more direct measure of movement kinematics. To carefully address this issue, we asked about the relationship between algorithm-identified behavioral transitions and behavioral changepoints identified by head-mounted inertial measurement units (IMUs), which allow us to capture precise 3D head orientation and linear acceleration while we record mice exploring an open field arena using an overhead 2D camera (Fig 4h). Behavioral transitions were identifiable in the IMU data as transient increases in the rates of change for acceleration (quantified by jerk) and orientation (quantified by angular velocity). Both measures correlated with state transitions identified by keypoint-MoSeq but failed to match transitions in behavioral states identified by B-SOiD, MotionMapper and VAME (Fig. 4i). Furthermore, IMU-extracted behavioral features (like head pitch or acceleration) typically rose and fell symmetrically around B-SOiD, MotionMapper and VAME-identified transitions, while keypoint-MoSeq identified asymmetrical changes in these features. For example, acceleration tended to be highest in the middle of B-SOiD-identified behavioral states, while acceleration tended to sharply change at the boundaries of keypoint-MoSeq-identified behavioral syllables (Fig 4i; Extended Data Fig 5a-b).

**Figure 5:**
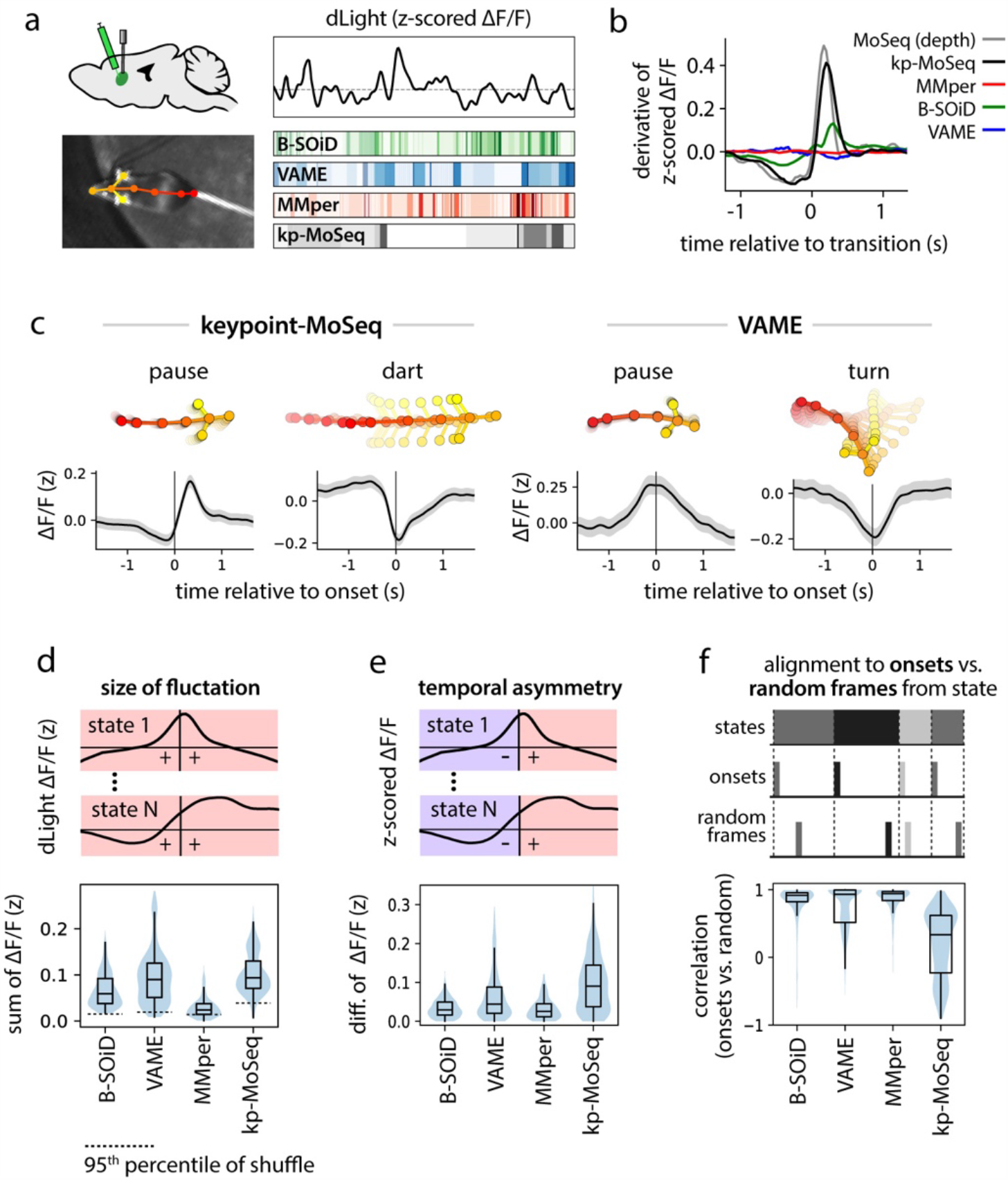
Keypoint-MoSeq syllable transitions align with fluctuations in striatal dopamine. **a)** Illustration depicting simultaneous recordings of dopamine fluctuations in dorsolateral striatum (DLS) obtained from fiber photometry (top) and unsupervised behavioral segmentation of 2D keypoint data (bottom). **b)** Derivative of the dopamine signal aligned to state transitions from each method. **c)** Average dopamine signal (z-scored ΔF/F) aligned to the onset of example states identified by keypoint-MoSeq and VAME. Shading marks the 95% confidence interval around the mean. **d)** Distributions capturing the magnitude of state-associated dopamine fluctuations across states from each method, where magnitude is defined as mean total absolute value in a one-second window centered on state onset. **e)** Distributions capturing the temporal asymmetry of state-associated dopamine fluctuations, where asymmetry is defined as the difference in mean dopamine signal during 500ms after versus 500ms before state onset. **f)** Temporal randomization affects keypoint-MoSeq identified neuro-behavioral correlations, but not those identified by other methods. **Top:** schematic of randomization. The dopamine signal was either aligned to the onsets of each state, as in (c), or to random frames throughout the execution of each state. **Bottom:** distributions capturing the correlation of state-associated dopamine fluctuations before vs. after randomization.

The fact that keypoint-MoSeq more clearly identifies behavioral boundaries does not necessarily mean that it is better at capturing the instantaneous content of behavior. Indeed, a spline-based linear encoding model was able to effectively reconstruct a panel of coarse kinematic parameters from all four of the explored methods with comparable accuracy (Extended Data Fig 4c). However, the fact that movement parameters – as measured by accelerometry – change suddenly at the onset of keypoint-MoSeq syllables, but not at the onset of B-SOiD, VAME or MotionMapper states, provide evidence that these methods afford fundamentally different views of temporal structure in behavior. The coincidence of behavioral transitions identified by keypoint-MoSeq (which are ultimately based on video data) and IMU data (which is based in movement per se) further validates the segmentation of behavior generated by keypoint-MoSeq.

### Keypoint-MoSeq state transitions align with fluctuations in neural data

Understanding the relationship between brain and behavior requires timestamps that enable researchers to align neural and behavioral data to moments of change. During traditional head-fixed behavioral tasks, such timestamps naturally arise out of task structure, in which time is divided up into clear, experimenter-specified epochs relating to e.g., the presentation of sensory cues or reward, the moment of behavioral report, etc. One of the main use cases for unsupervised behavioral classification is to understand how the brain generates spontaneous behaviors that arise outside of a rigid task structure^9^; in this setting, the boundaries between behavioral states serve as surrogate timestamps to allow alignment of neural data.

We have recently used depth MoSeq to show that the levels of the neuromodulator dopamine fluctuate within the dorsolateral striatum (DLS) during spontaneous behavior, and that these fluctuations are temporally aligned to syllable transitions^18^: On average, dopamine levels rise rapidly at the onset of each syllable, and then decline toward the end of the syllable. Furthermore, the average magnitude of dopamine fluctuations varies across syllables. We wondered whether we could recapitulate these previously observed relationships between syllable transitions and dopamine fluctuations using keypoint-MoSeq or alternative methods for fractionating keypoint data into behavioral states (Fig 5a).

Syllable-associated dopamine fluctuations (as captured by dLight photometry) were remarkably similar between depth MoSeq and keypoint-MoSeq; aligning the derivative of the dopamine signal to syllable transitions revealed a trajectory that was almost identical between depth MoSeq and keypoint-MoSeq, with a shallow dip prior to syllable onset and sharp rise after onset (Fig 5b). State-related dopamine fluctuations were much lower in amplitude (or non-existent), however, when assessed using B-SOiD, VAME and MotionMapper (Fig 5b). Given the association between striatal dopamine release and movement^28^, it is possible that method-to-method variation can be explained by differences in how each method represents stationary vs. locomotory behavior. Yet, the transition-associated dopamine fluctuations highlighted by keypoint-MoSeq remained much more prominent than those from other methods when analysis was restricted to high or low velocity states (Extended Data Fig 6a).

**Figure 6:**
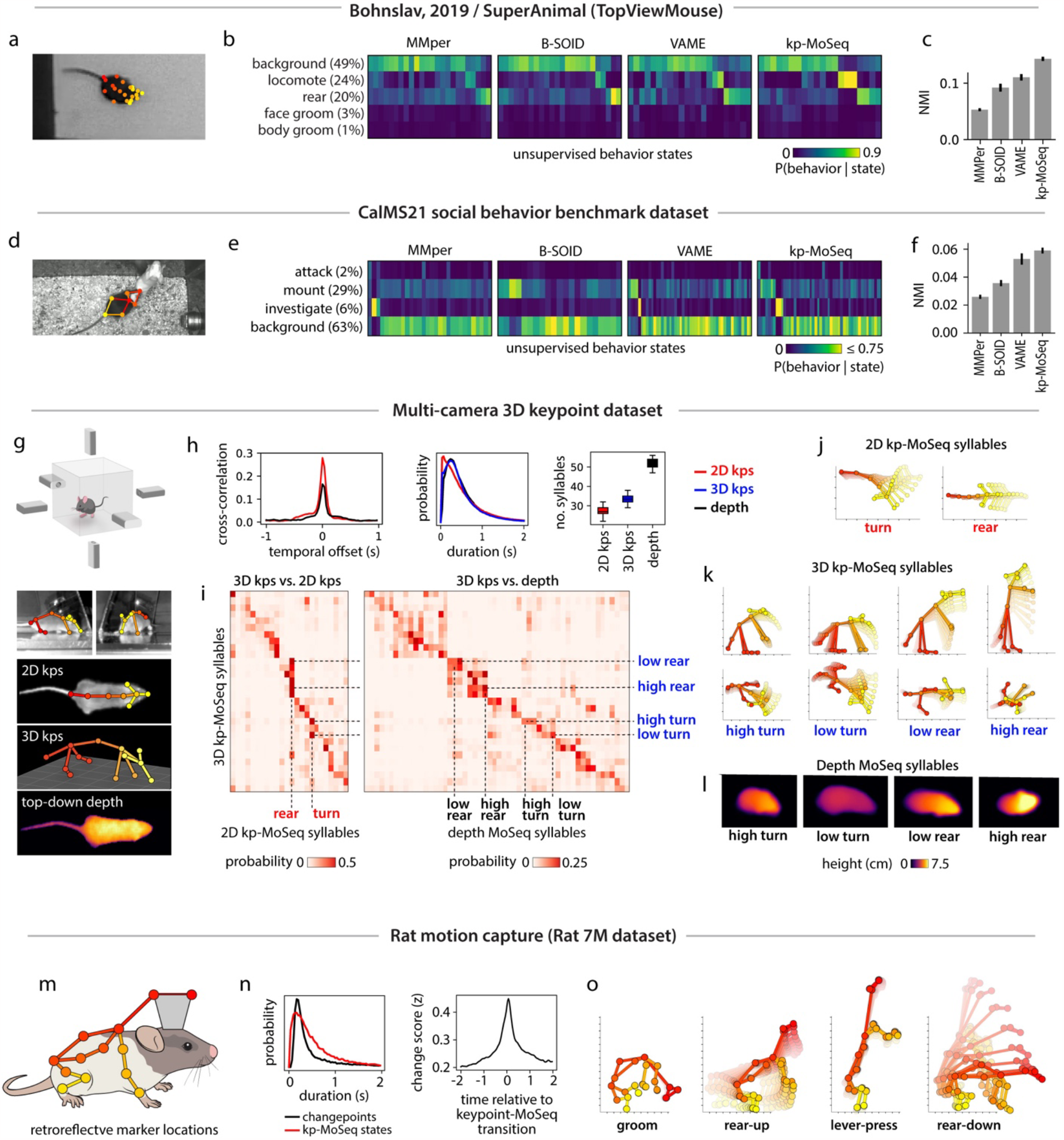
Keypoint-MoSeq generalizes across pose representations, behaviors, and rodent species. **a)** Example frame from a benchmark open field dataset (Bohnslav, 2019). **b)** Overall frequency of each human-annotated behavior (as %) and conditional frequencies across states inferred from unsupervised analysis of 2D keypoints. **c)** Normalized mutual information (NMI, see Methods) between human annotations and unsupervised behavior labels from each method. **d)** Example frame from the CalMS21 social behavior benchmark dataset, showing 2D keypoint annotations for the resident mouse. **e-f)** Overlap between human annotations and unsupervised behavior states inferred from 2D keypoint tracking of the resident mouse, as b-c. **g)** Multi-camera arena for simultaneous recording of 3D keypoints (3D kps), 2D keypoints (2D kps) and depth videos. **h)** Comparison of model outputs across tracking modalities. 2D and 3D keypoint data were modeled using keypoint-MoSeq, and depth data were modeled using original MoSeq. **Left:** cross-correlation of transition rates, comparing 3D keypoints to 2D keypoints and depth respectively. **Middle:** distribution of syllable durations; **Right:** number of states with frequency > 0.5%. Boxplots represent the distribution of state counts across 20 independent runs of each model. **i)** Probability of syllables inferred from 2D keypoints (left) or depth (right) during each 3D keypoint-based syllable. **j-l)** Average pose trajectories for the syllables marked in (i). **k)** 3D trajectories are plotted in side view (first row) and top-down view (second row). **l)** Average pose (as depth image) 100ms after syllable onset. **m)** Location of markers for rat motion capture. **n) Left:** Average keypoint change score (z) aligned to keypoint-MoSeq transitions. **Right:** Duration distributions for keypoint-MoSeq states and inter-changepoint intervals. **o)** Average pose trajectories for example syllables learned from rat motion capture data.

We wondered whether the inability of alternative clustering methods to identify a clear relationship between spontaneous behavior and dopamine could be explained by differences in how they represent the temporal structure of behavior. If, as we have shown, B-SOiD, VAME and MotionMapper can capture the content of behavior but not the timing of transitions, then one might expect average dopamine levels to vary consistently across the different behavioral states identified by these alternative methods. To test this prediction, we computed the average dopamine trace aligned to state onset separately for each state (Fig 5c). Across all methods almost every state was associated with a consistent average increase or decrease in dopamine levels (Fig 5c-d, Extended Data Fig 6b).

However, the specific pattern of fluctuation identified by each method substantially varied. Dopamine tended to increase at the initiation of keypoint-MoSeq-identified behavioral syllables, with dopamine baselines and amplitudes varying across syllables. In contrast, dopamine signals were typically at a peak or nadir at the beginning of each state identified by alternative methods, forming a trajectory that was symmetric around state onset (Fig 5c). This symmetry tended to wash out dopamine dynamics, with the average change in the dopamine signal before vs. after syllable onset being approximately three times larger for keypoint-MoSeq than for alternative methods (Fig 5e). Similarly, the number of states where the z-scored dopamine signal changed sign before vs. after state onset was ∼2-fold greater for keypoint-MoSeq than for alternatives. Furthermore, aligning the dopamine signal to randomly-sampled times throughout the execution of each behavioral state – rather than its onset – radically altered the state-associated dopamine dynamics observed using keypoint-MoSeq, but made little difference for alternative methods (Fig 5f, Extended Data Fig 6c-d), a result that could not be explained simply by differences in each state’s duration (Extended Data Fig 6c). These results suggest that the onsets of keypoint-MoSeq-identified behavioral syllables are meaningful landmarks for neural data analysis, while state onsets identified by alternative methods are often functionally indistinguishable from timepoints randomly chosen from throughout the duration of a behavior.

### Keypoint-MoSeq generalizes across pose representations and behaviors

Keypoint tracking is a powerful means of pose estimation because it is so general: available methods can be flexibly applied to a wide variety of experimental setups, can capture diverse behaviors, and afford the experimenter broad latitude in the choice of which parts to track and at what resolution. To test the ability of keypoint-MoSeq to generalize across laboratories — and to better understand the mapping between syllables and human-identified behaviors — we used keypoint-MoSeq and alternative methods to analyze a pair of published benchmark datasets^29,30^. The first dataset included conventional 2D videos of a single mouse behaving in an open field, with human annotations for four commonly occurring behaviors (locomote, rear, face groom and body groom) (Fig 6a-c). To identify keypoints in this dataset we used DeepLabCut, specifically the TopViewMouse SuperAnimal network from the DLC Model Zoo^31^, which automatically identifies keypoints without the need for annotation data or training. The second dataset (part of the CalMS21 benchmark^30^) included a set of three manually annotated social behaviors (mounting, investigation, and attack) as well as keypoints for a pair of interacting mice (Fig 6d-f).

Changepoints analysis of keypoint data from both datasets identified block-like structure whose mean duration was ∼400ms, consistent with the presence of a behavioral rhythm organized at the sub-second timescale (Extended Data Fig 7a-b). Consistent with this, Keypoint-MoSeq recovered syllables from both datasets whose average duration was ∼400ms while, as before, the B-SOiD, MotionMapper and VAME identified behavioral states that were much shorter (Extended Data Fig 7c-d). Keypoint-MoSeq was also better at conveying information about which human-identified behavioral states were occurring at each moment than alternative methods; that said, the different methods were not dramatically different in terms of quantitative performance, consistent with each doing a reasonable job of capturing broad information about behavior (Fig 6c,f, Extended Data Fig 7e-f). However, there were some important differences: in the CalMS21 dataset, for example, MotionMapper, B-SOiD and VAME only identified a single behavior consistently (by defining a state specific to that behavior); B-SOiD and VAME only captured mounting and MotionMapper only captured investigation in 100% of model fits. Keypoint-MoSeq, in contrast, defined at least one state specific to each of the three behaviors in 100% of model fits (Extended Data Fig 7g). These results demonstrate that keypoint-MoSeq can identify temporal structure in diverse 2D keypoint datasets and reveal consistency between keypoint-MoSeq and supervised labels for behavioral states.

The above benchmark datasets differ widely in the number of keypoints tracked (7 for CalMS21 vs. 21 for the TopViewMouse model; Fig 6a,d), raising the question of how the pose representation fed to keypoint-MoSeq influences its outputs. Comparing keypoints to depth offers one clue: we noted that the number of syllables (∼25) identified in our open field data by keypoint-MoSeq using 2D keypoints was substantially fewer than the number identified by depth MoSeq (∼50). These findings suggest that higher dimensional input data – such as depth – affords MoSeq more information about pose during spontaneous behavior, which in turn yields a richer behavioral description. To test this hypothesis rigorously, we used multiple cameras to estimate the position of keypoints in 3D (including 6 keypoints that were not visible in the overhead camera 2D dataset) (Fig 6g). Compared to the 2D data, the new 3D keypoint pose representation was higher dimensional, had smoother trajectories and exhibited oscillatory dynamics related to gait (Extended Data Fig 8a-b). Yet the temporal structure of both the data and the syllables that emerged from keypoint-MoSeq was surprisingly similar: the 3D data contained similar changepoints to both the 2D and depth data (Extended Data 8c-d), and after processing with keypoint-MoSeq the resulting syllable duration distributions were almost identical between the 2D, 3D and depth datasets, and syllable transitions tended to occur at the same moments in time (Fig 6h).

There was a bigger change, however, in the way syllables were categorized when comparing 2D and 3D data. Keypoint-MoSeq tended to distinguish more syllable states in the 3D data (33±2 syllables for 3D keypoints vs. 27±2 syllables for 2D keypoints and 52±3 syllables for depth MoSeq; Fig 6h, Suppl Movie 4), especially for behaviors in which height varied (Fig 6i). Turning, for example, was grouped as a single state with the 2D keypoint data but partitioned into three states with different head positions with the 3D keypoint data (nose to the ground vs. nose in the air), and seven different states in the depth data (Fig 6j-l). Rearing was even more fractionated, with a single 2D syllable splitting six ways based on body angle and trajectory in the 3D keypoint data (rising vs. falling) and 8 ways in the depth data. These data demonstrate that keypoint-MoSeq works well on both 2D and 3D keypoint data; furthermore, our analyses suggest that higher-dimensional sources of input data to MoSeq give rise to richer descriptions of behavior, but that even relatively low-dimensional 2D keypoint data can be used to usefully identify behavioral transitions.

Finally, to test if keypoint-MoSeq generalizes across species, we analyzed previously published 3D motion capture data derived from rats. In this dataset, rats were adorned with reflective body piercings and recorded in a circular home cage arena with a lever and water spout for operant training (Fig 6m; Rat7M dataset^32^). As with mice, changepoint analysis identified sub-second blocks of continuous kinematics (Fig 6n; Extended Data Fig 9a). Keypoint-MoSeq captured this temporal structure, and identified syllables whose transitions that aligned with changepoints in the keypoint data (Fig 6n). As was true in the mouse data, rat syllables included a diversity of behaviors, including a syllable specific to lever-pressing in the arena (Fig 6o; Extended Data Fig 9b; Suppl Movie 5).

## Discussion

MoSeq is a well-validated method for behavioral segmentation that leverages natural sub-second discontinuities in rodent behavior to automatically identify the behavioral syllables out of which spontaneous behavior is assembled^17-20^. However, the conventional MoSeq platform is unable to directly accept keypoint data, as pervasive keypoint jitter (a previously-characterized limitation of neural network-based pose tracking^5,22^) causes MoSeq to identify false behavioral transitions^13,22^. To address this challenge, here we describe keypoint-MoSeq, an SLDS model that enables joint inference of keypoint positions and associated behavioral syllables. Keypoint-MoSeq effectively estimates syllable structure in a wide variety of circumstances (e.g., in mice or rats, in video shot from above or below, in data capturing 2D or 3D keypoints, in animals behaving alone or during social interactions, in mice with or without headgear or neural implants). We validate keypoint-MoSeq by demonstrating that identified behavioral syllables are interpretable; that their transitions match changepoints in depth and kinematic data; and that the syllables capture systematic fluctuations in neural activity and complex behaviors identified by expert observers. Thus keypoint-MoSeq affords much of the same insight into behavioral structure as depth MoSeq, while rendering behavioral syllables and grammar accessible to researchers who use standard video to capture animal behavior.

There are now many techniques for unsupervised behavior segmentation^9,33^. The common form of their outputs – a sequence of discrete labels – belies profound variation in how they work and the kinds of biological insight one might gain from applying them. To better understand their relative strengths and weaknesses when applied to mouse keypoint data, here we perform a detailed head-to-head comparison between keypoint-MoSeq and three alternative methods (B-SOiD^12^, MotionMapper^23^ and VAME^13^). All these methods similarly encode the kinematic content of mouse behavior. The methods differed radically, however, in the temporal structure of their outputs. Keypoint-MoSeq syllables lasted almost an order of magnitude longer on average than states identified by alternative clustering methods, and transitions between B-SOiD, MotionMapper and VAME states often occurred in the middle of what a human might identify as a behavioral module or motif (e.g., a rear). Our analysis suggests two possible reasons for this difference. First, unlike alternative methods, MoSeq can discretize behavior at a particular user-defined timescale, and therefore is better able to identify clear boundaries between behavioral elements that respect the natural sub-second rhythmicity in mouse movement and neural activity. The resulting parsimony prevents over-fractionation of individuals behaviors, as we observed when clustering keypoint data using alternative methods. Second, the hierarchical structure of keypoint-MoSeq’s underlying generative model means it can detect noise in keypoint noise trajectories and distinguish this noise from actual behavior without smoothing away meaningful behavioral transitions.

The fact that MoSeq is a probabilistic generative model means that its descriptions of behavior are constrained by the model structure and its parameters: it seeks to describe behavior as composed of auto-regressive trajectories through a pose space with switching dynamics organized at a single main timescale. Because MoSeq instantiates an explicit model for behavior, there are many tasks in behavioral analysis for which keypoint-MoSeq may be ill-suited. For example, as has been previously noted, keypoint-MoSeq cannot integrate dynamics across a wide range of timescales, as would be possible with methods such as MotionMapper^34,35^. In addition, some behaviors — like the leg movements of walking flies — may be better captured by methods whose design emphasizes oscillatory dynamics. It is important to note that, despite its structural constraints, MoSeq is not *only* useful for capturing fine timescale structure in behavior; indeed, MoSeq has repeatedly been shown to be performant at tasks that pervasively influence the structure of behavior, including changes in behavior due to genetic mutations or drug treatments^17,20^. That said, we stress that there is no one “best” approach for behavioral analysis, as all methods involve trade-offs: methods that work for one problem (for example, identifying fast neurobehavioral correlates) may not be well suited for another problem.

The outputs of MoSeq depend upon the type of data it is fed. While similar behavioral boundaries are identified from 2D keypoints, 3D keypoints and depth data, increasing the dimensionality of the input data also increases the richness of the syllables revealed by MoSeq. Though directly modeling the raw pixel intensities of depth^17^ or 2D video^36^ recordings provides the most detailed access to spontaneous behavior, technical challenges (ranging from reflection sensitivity to relatively low temporal resolution) can make depth cameras difficult to use in many experimental settings. Similarly, occlusions and variation in perspective and illumination remain a challenge for direct 2D video modeling. The development of keypoint-MoSeq — together with the extraordinary advances in markerless pose tracking — should enable MoSeq to be used in a variety of these adversarial circumstances, such as when mice are obstructed from a single axis of view, or when the environment changes dynamically. In addition, keypoint-MoSeq can also be applied to the petabytes of legacy data sitting fallow on the hard drives of investigators who have already done painstaking behavioral experiments using conventional video cameras. Going forward, increasingly sophisticated pose tracking approaches^22,37^ and methods that combine keypoint tracking with direct video analysis^38^ may eventually close the gap in dimensionality between keypoint- and (depth) video-based pose tracking.

To facilitate the adoption of keypoint-MoSeq we have built a website (www.MoSeq4all.org) that includes free access to the code for academics as well as extensive documentation and guidance for implementation. As demonstrated by this paper, the model underlying MoSeq is modular and therefore accessible to extensions and modifications that can increase its alignment to behavioral data. For example, Costacurta et al., recently reported a time-warped version of MoSeq that incorporates a term to explicitly model variation in movement vigor^39^. We anticipate that the application of keypoint-MoSeq to a wide variety of experimental datasets will both yield important information about the strengths and failure modes of model-based methods for behavioral classification, and prompt continued innovation.

## Acknowledgements

S.R.D. is supported by NIH grants RF1AG073625, R01NS114020, U24NS109520, the Simons Foundation Autism Research Initiative, and the Simons Collaboration on Plasticity and the Aging Brain. S.R.D. and S.W.L are supported by NIH grant U19NS113201 and the Simons Collaboration on the Global Brain. C.W. is a Fellow of the Jane Coffin Childs Memorial Fund for Medical Research. W.F.G. is supported by NIH grant F31NS113385. M.J. is supported by NIH grant F31NS122155. S.W.L is supported by the Alfred P. Sloan Foundation. T.P. is supported by a Salk Collaboration Grant. We thank J. Araki for administrative support; the HMS Research Instrumentation Core, which is supported by the Bertarelli Program in Translational Neuroscience and Neuroengineering, and by NEI grant EY012196; and members of the Datta laboratory for useful comments on the paper. Portions of this research were conducted on the O2 High Performance Compute Cluster at Harvard Medical School.

## Code availability

Software links and user-support for both depth and keypoint data are available at the MoSeq homepage: MoSeq4all.org. Data loading, project configuration and visualization are enabled through the “keypoint-moseq” python library (https://github.com/dattalab/keypoint-moseq). We also developed a standalone library called “jax-moseq” for core model inference (https://github.com/dattalab/jax-moseq). Both libraries are freely available to the research community.

## EXPERIMENTAL METHODS

### Animal care and behavioral experiments

Unless otherwise noted, behavioral recordings were performed on 8–16-week-old C57/BL6 mice (The Jackson Laboratory stock no. 000664). Mice were transferred to our colony at 6-8 weeks of age and housed in a reverse 12-hour light/12-hour dark cycle. We single-housed mice after stereotactic surgery, and group-housed them otherwise. On recording days, mice were brought to the laboratory, habituated in darkness for at least 20 minutes, and then placed in an open field arena for 30-60 mins. We recorded 6 male mice for 10 sessions (6 hours) in the initial round of open field recordings; and 5 male mice for 52 sessions (50 hours) during the accelerometry recordings. The dopamine photometry recordings were obtained from a recent study^1^. They include 6 C57/BL6 mice and 8 DAT-IRES-cre (The Jackson Laboratory stock no. 006660) mice of both sexes, recorded for 378 sessions. Of these, we selected a random subset of 95 sessions (∼50 hours) for benchmarking keypoint-MoSeq.

### Stereotactic surgery procedures

For all stereotactic surgeries, mice were anaesthetized using 1–2% isoflurane in oxygen, at a flow rate of 1 L/min for the duration of the procedure. Anterior-posterior (AP) and medial-lateral (ML) coordinates were zeroed relative to bregma, the dorsoventral (DV) coordinate was zeroed relative to the pial surface, and coordinates are in units of mm. For dopamine recordings, 400nL of AAV5.CAG.dLight1.1 (Addgene #111067, titer: 4.85 × 10^12^) was injected at a 1:2 dilution into the DLS (AP 0.260; ML 2.550; DV −2.40) and a single 200-μm diameter, 0.37–0.57 NA fiber cannula was implanted 200 μm above the injection site (see ref^1^ for additional details). For accelerometry recordings, we surgically attached a millmax connector (DigiKey ED8450-ND) and head bar to the skull and secured it with dental cement (Metabond). A 9 degree-of-freedom absolute orientation inertial measurement unit (IMU; Bosch BN0055) was mounted on the millmax connector using a custom printed circuit board (PCB) with a net weight below 1g.

### Data acquisition from the IMU

The IMU was connected to a Teensy microcontroller, which was programmed using the Adafruit BNO055 library with default settings (sample rate: 100 Hz, units: m/s^2^). To synchronize the IMU measurements and video recordings, we used an array of near infrared LEDs to display a rapid sequence of random 4-bit codes that updated throughout the recording. The code sequence was later extracted from the behavioral videos and used to fit a piecewise linear model between timestamps from the videos and timestamps from the IMU.

### Recording setup

For the initial set of open field recordings (Fig 1-3, 4a-g Fig 6g-l), mice were recorded in a square arena with transparent floor and walls (30cm length and width). Microsoft Azure Kinect cameras captured simultaneous depth and near-infrared video at 30Hz. Six cameras were used in total: one above, one below, and four side cameras at right angles at the same height as the mouse. For the accelerometry recordings, we used a single Microsoft Azure Kinect camera placed above the mouse, and an arena with transparent floor and opaque circular walls (45cm diameter). Data was transferred from the IMU using a light-weight tether attached to a custom-built active commutator. For the dopamine perturbation experiments, we used a slightly older camera model – the Microsoft Kinect 2 – to capture simultaneous depth and near-infrared at 30Hz. The recording arena was circular with opaque floor and walls (45cm diameter). Photometry signals were conveyed from the mouse using a fiber-optic patch cord attached to a passive commutator.

## COMPUTATIONAL METHODS

### Processing depth videos

Applying MoSeq to depth videos involves: (1) mouse tracking and background subtraction; (2) egocentric alignment and cropping; (3) principal component analysis (PCA); (4) probabilistic modeling. We applied steps (2-4) as described in the MoSeq2 pipeline^2^. For step (1), we trained a convolutional neural network (CNN) with a Unet++^3^ architecture to segment mouse from background using ∼5000 hand-labeled frames as training data.

### Keypoint tracking

We used CNNs with an HRNet^4^ architecture (https://github.com/stefanopini/simple-HRNet) with a final stride of 2 for pose tracking. The networks were trained on ∼1000 hand-labeled frames each for the overhead, below-floor, and side-view camera angles. Frame-labelling was crowdsourced through a commercial service (Scale AI). For the overhead camera, we tracked two ears and 6 points along the dorsal midline (tail base, lumbar spine, thoracic spine, cervical spine, head, and nose). For the below-floor camera, we tracked the tip of each forepaw, the tip and base of each hind paw, and four points along the ventral midline (tail base, genitals, abdomen, and nose). For the side cameras, we tracked the same eight points as for the overhead camera, and also included the six limb points that were used for the below-floor camera (14 total). We trained a separate CNN for each camera angle. Target activations were formed by centering a Gaussian with 10px standard deviation on each keypoint. We used the location of the maximum pixel in each output channel of the neural network to determine keypoint coordinates and used the value at that pixel to set the confidence score. The resulting mean absolute error (MEA) between network detections and manual annotations was 2.9 pixels (px) for the training data and 3.2 px for heldout data. We also trained DeepLabCut and SLEAP models on the overhead-camera and below-floor-camera datasets. For DeepLabCut, we used version 2.2.1, setting the architecture to resnet50 architecture and the “pos_dist_thresh” parameter to 10, resulting in train and test MEAs of 3.4 px and 3.8 px respectively. For SLEAP, we used version 1.2.3 with the baseline_large_rf.single.json configuration, resulting in train and test MEAs of 3.5 px and 4.7 px.

### 3D pose inference

Using 2D keypoint detections from six cameras, 3D keypoint coordinates were triangulated and then refined using GIMBAL, a model-based approach that leverages anatomical constraints and motion continuity^5^. To fit GIMBAL, we computed initial 3D keypoint estimates using robust triangulation (i.e. by taking the median across all camera pairs, as in 3D-DeepLabCut^6^) and then filtered to remove outliers using the EllipticEnvelope method from sklearn; We then fit the skeletal parameters and directional priors for GIMBAL using expectation maximization with 50 pose states (see ref^5^ for details). Finally, we applied the fitted GIMBAL model to each recording, using the following parameters for all keypoints: obs_outlier_variance=1e6, obs_inlier_variance=10, pos_dt_variance=10. The latter parameters were chosen based on the accuracy of the resulting 3D keypoint estimates, as assessed from visual inspection.

### Inferring model-free changepoints

We defined changepoints as sudden, simultaneous shifts in the trajectories of multiple keypoints. We detected them using a procedure similar to the filtered derivative algorithm described in ref^7^, but with changes to emphasize simultaneity across multiple keypoints. The changes account for the lower dimensionality of keypoint data compared to depth videos, and for the unique noise structure of markerless keypoint tracking, in which individual keypoints occasionally jump a relatively large distance due to detection errors. Briefly, the new procedure first defines a continuous change score by: (1) calculating the rate of each in each keypoint coordinate; (2) quantifying simultaneity in the change-rates across keypoints; (3) transforming the signal based on statistical significance with respect to a temporally shuffled null distribution; (4) identifying local peaks in the resulting significance score. The details of each step are as follows.

1. **Calculating rates of change:** We transformed the keypoint coordinates on each frame by centering and aligned them along the tail-nose axis. We then computed the derivative of each coordinate for each keypoint, using a sliding window of length 3 as shown below, where *x*_*t*_ denotes the value of a coordinate at time *t*.

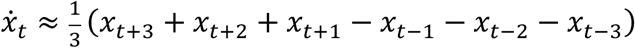
2. **Quantifying simultaneous changes:** The derivatives for each keypoint were Z-scored and then binarized with a threshold. We then counted the number of threshold crossings on each frame and smoothed the resulting time-series of counts using a Gaussian filter with a one-frame kernel. The value of the threshold was chosen to maximize the total number of detected changepoints.
3. **Comparing to a null distribution:** We repeated step (2) for 1000 shuffled datasets, in which each keypoint trajectory was cyclically permuted by a random interval. Using the shuffles as a null distribution, we computed a P-value for each frame and defined the final change score as – log_10_(pval)
4. **Identifying local peaks in the change score:** We identified local peaks in the change score *s*_*t*_, i.e., times *t* for which *s*_*t*−1_ < *s*_*t*_ > *s*_*t*+1_. Peaks were classified as statistically significant when they corresponded to a p-value below 0.01, which was chosen to control the false-discovery rate at 10%. The statistically significant peaks were reported as changepoints for downstream analysis.

### Spectral Analysis

To analyze keypoint jitter, we quantified the magnitude of fluctuations across a range of frequencies by computing a spectrogram for each keypoint along each coordinate axis. Spectrograms were computed using the python function scipy.signal.spectrogram with nperseg=128 and noverlap=124. The spectrograms were then combined through averaging: each keypoint was assigned a spectrogram by averaging over the two coordinate axes, and the entire animal was assigned a spectrogram by averaging over all keypoints.

We used the keypoint-specific spectrograms to calculate cross-correlations with −log_10_(neural network detection confidence), as well as the “error magnitude” (Fig 2f). Error magnitude was defined as the distance between the detected 2D location of a keypoint (based on a single camera angle) and a reprojection of its 3D position (based on consensus across six camera angles; see “3D pose inference” above). We also computed the cross-correlation between nose- and tail-base-fluctuations at each frequency, as measured by the overhead and below-floor cameras respectively. Finally, we averaged spectral power across keypoints to compute the cross-correlation with model transition rates (Fig 2f), defined as the per-frame probability of a state transitions across 20 model restarts.

### Applying keypoint-MoSeq

The initial open field recordings (Fig 1-4), as well as the accelerometry, dopamine, and two benchmark datasets were modeled separately. Twenty models with different random seeds were fit for each dataset (except for the accelerometry data, in which case one model was fit).

Modeling consisted of two phases: (1) Fitting an autoregressive hidden Markov model (AR-HMM) to a fixed pose trajectory derived from PCA of egocentric-aligned keypoints; (2) Fitting a full keypoint-MoSeq model initialized from the AR-HMM. References in the text to “MoSeq applied to keypoints” or “MoSeq (keypoints)”, e.g., in Figs 2-3, refer to output of step (1). Both steps are described below, followed by a detailed description of the model and inference algorithm in the mathematical modeling section. In all cases, we excluded rare states (frequency < 0.5%) from downstream analysis. We have made the code available as a user-friendly package, available at Moseq4all.org.

1. Fitting an initial AR-HMM: We first modified the keypoint coordinates, defining keypoints with confidence below 0.5 as missing data and in imputing their values via linear interpolation, and then augmenting all coordinates with a small amount of random noise; The noise values were uniformly sampled from the interval [-0.1, 0.1] and helped prevent degeneracy during model fitting. Importantly, these preprocessing steps were only applied during AR-HMM fitting – the original coordinates were used when fitting the full keypoint-MoSeq model. Next, we centered the coordinates on each frame, aligned them using the tail-nose angle, and then transformed them using PCA with whitening. The number of principal components (PCs) was chosen for each dataset as the minimum required to explain 90% of total variance. This resulted in 4 PCs for the overhead camera 2D datasets, 6 PCs for the below-floor-camera 2D datasets, and 6 PCs for the 3D dataset. We then used Gibbs sampling to infer the states and parameters of an AR-HMM, including the state sequence *z*, the autoregressive parameters *A, b, Q*, and the transition parameters *π, β*. The hyper-parameters for this step, listed in the mathematical modeling section below, were generally identical to those in the original depth-MoSeq model^7^. The one exception was κ which we adjusted separately for each dataset to ensure a median state duration of 400ms.
2. Fitting a full keypoint-MoSeq model: We next fit the full set of variables for keypoint-MoSeq, which include the AR-HMM variables mentioned above, as well as the location *v* and heading *h*, latent pose trajectory *x*, per-keypoint noise level σ^2^, and per-frame/per-keypoint noise scale *s*. Fitting was performed using Gibbs sampling for 500 iterations, at which point the log joint probability appeared to have stabilized. The hyper-parameters for this step are enumerated in the mathematical modeling section below. In general, we used the same hyper-parameter values across datasets. The two exceptions were *κ*, which again had to be adjusted to maintain a median state duration of 400ms, and *s*_0_, which determines a prior on the noise scale. Since low-confidence keypoint detections often have high error, we set *s*_0_ using a logistic curve that transitions between a high-noise regime (*s*_0_ = 100) for detections with low confidence and a low-noise regime (*s*_0_ = 1) for detections with high confidence:

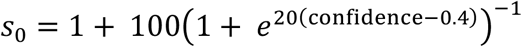

### Trajectory plots

To visualize the modal trajectory associated with each syllable (Fig 3e), we (1) computed the full set of trajectories for all instances of all syllables (2) used a local density criterion to identify a single representative instance of each syllable (3) computed a final trajectory using the nearest neighbors of the representative trajectory.

1. Computing the trajectory of individual syllable instances: Let *y*_*t*_, *v*_*t*_, and *h*_*t*_ denote the keypoint coordinates, centroid and heading of the mouse at time *t*, and let *F(v, h; y*) denote the rigid transformation that egocentrically aligns *y* using centroid *v* and heading *h*. Given a syllable instance with onset time *T*, we computed the corresponding trajectory *X*_*T*_ by centering and aligning the sequence of poses (*y*_*T*−5_, …, *y*_*T*+15_) using the centroid and heading on time *T*. In other words,

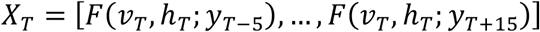
2. Identifying a representative instance of each syllable: The collection of trajectories computed above can be thought of as a set of points in a high dimensional trajectory space (for *K* keypoints in 2D, this space would have dimension 40*K*). Each point has a syllable label, and the segregation of these labels in the trajectory space represents the kinematic differences between syllables. To capture these differences, we computed a local probability density function for each syllable, and a global density function across all syllables. We then selected a representative trajectory *X* for each syllable by maximizing the ratio:

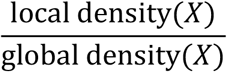

The density functions were computed as the mean distance from each point to its 50 nearest neighbors. For the global density, the nearest neighbors were selected from among all instances of all syllables. For the local densities, the nearest neighbors were selected from among instances of the target syllable.
3. Computing final trajectories for each syllable: For each syllable and its representative trajectory *X*, we identified the 50 nearest neighbors of *S* from among other instanes of the same syllable and then computed a final trajectory as the mean across these nearest neighbors. The trajectory plots in Fig 3e consist of 10 evenly-space poses along this trajectory, i.e., the poses at times *T* − 5, *T* − *3*, …, *T* + 1*3*.

### Cross-syllable likelihoods

We defined each cross-syllable likelihood^7^ as the probability (on average) that instances of one syllable could have arisen based on the dynamics of another syllable. The probabilities were computed based on the discrete latent states *z*_*t*_, continuous latent states *x*_*t*_, and autoregressive parameters *A, b, Q* output by keypoint-MoSeq. The instances *I(n*) of syllable *n* were defined as the set of all sequences (*t*_*s*_, …, *t*_*e*_) of consecutive timepoints such that *z*_*t*_ = *n* for all *t*_*s*_ ≤ *t* ≤ *t*_*e*_ and 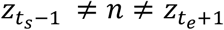. For each such instance, one can calculate the probability 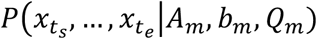 that the corresponding sequence of latent states arose from the autoregressive dynamics of syllable *m*. The cross-syllable likelihood *C*_*nm*_ is defined in terms of these probabilities as

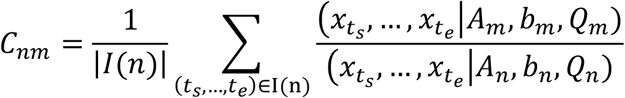

### Generating synthetic keypoint data

To generate the synthetic keypoint trajectories used for Extended Data Fig 3c, we fit a linear dynamical system (LDS) to egocentrically aligned keypoint trajectories and then sampled randomly generated outputs from the fitted model. The LDS was identical to the model underlying keypoint-MoSeq (see mathematical modeling section below), except that it only had one discrete state, lacked centroid ad heading variables, and allowed separate noise terms for the x- and y-coordinates of each keypoint.

### Applying B-SOiD

B-SOiD is an automated pipeline for behavioral clustering that: (1) preprocesses keypoint trajectories to generate pose and movement features; (2) performs dimensionality reduction on a subset of frames using UMAP; (3) clusters points in the UMAP space; (4) uses a classifier to extend the clustering to all frames^8^. We fit B-SOiD separately for each dataset. In each case, steps 2-4 were performed 20 times with different random seeds, and the pipeline was applied with standard parameters; 50,000 randomly sampled frames were used for dimensionality reduction and clustering, and the min_cluster_size range was set to 0.5% - 1%. Since B-SOiD uses a hardcoded window of 100ms to calculate pose and movement features, we re-ran the pipeline with falsely inflated framerates for the window-size scan in Extended Data Fig 4a. In all analyses involving B-SOiD, rare states (frequency < 0.5%) were excluded from analysis.

### Applying VAME

VAME is a pipeline for behavioral clustering that: (1) preprocesses keypoint trajectories and transforms them into egocentric coordinates; (2) fits a recurrent neural network (RNN); (3) clusters the latent code of the RNN^9^. We applied these steps separately to each dataset, in each case running step (3) 20 times with different random seeds. For step (1), we used the same parameters as in keypoint-MoSeq – egocentric alignment was performed along the tail-nose axis, and we set the pose_confidence threshold to 0.5. For step (2), we set time_window=30 and zdims=20 for all datasets, except for the zdim-scan in Extended Data Fig 4a. VAME provides two different options for step (3): fitting an HMM (default) or applying K-Means (alternative). We fit an HMM for all datasets and additionally applied K-Means to the initial open dataset. In general, we approximately matched the number of states/clusters in VAME to the number identified by keypoint-MoSeq, except when scanning over state number in Extended Data Fig 4a. In all analyses involving VAME, rare states (frequency < 0.5%) were excluded from analysis.

### Applying MotionMapper

MotionMapper performs unsupervised behavioral segmentation by: (1) applying a wavelet transform to preprocessed pose data; (2) nonlinearly embedding the transformed data in 2D; (3) clustering the 2D data with a watershed transform^10^. We applied MotionMapper separately to each dataset using the python package https://github.com/bermanlabemory/motionmapperpy. In general, the data were egocentrically aligned along the tail-nose axis and then projected into 8 dimensions using PCA. 10 log-spaced frequencies between 0.25 and 15Hz were used for the wavelet transform, and dimensionality reduction was performed using tSNE. The threshold for watershedding was chosen to produce at least 25 clusters, consistent with keypoint-MoSeq for the overhead camera data. Rare states (frequency < 0.5%) were excluded from analysis. For the parameter scan in Extended Data Fig 4a, we varied the each of these parameters while holding the others fixed, including the threshold for watershedding, the number of initial PCA dimensions, and the frequency range of wavelet analysis. We also repeated a subset of these analyses using an alternative autoencoder-based dimensionality reduction approach, as described in the motionmapperpy tutorial (motionmapperpy/demo/motionmapperpy_mouse_demo.ipynb).

### Predicting kinematics from state sequences

We trained decoding models based on spline regression to predict kinematic parameters (height, velocity, turn speed) from state sequences output by keypoint-MoSeq and other behavior segmentation methods (Fig 4e, Extended Data Fig 4c). Let *z*_*t*_ represent an unsupervised behavioral state sequence and let *B* denote a spline basis, where *B*_*t,i*_ is the value of spline *i* and frame *t*. We generated such a basis using the “bs” function from the python package “patsy”, passing in five log-spaced knot locations (1.0, 2.0, 3.9, 7.7, 15.2, 30.0) and obtaining basis values over a 300-frame interval. This resulted in a 300-by-5 basis matrix *h*. The spline basis and state sequence were combined to form a 5*N*-dimensional design matrix, where *N* is the number of distinct behavioral states. Specifically, for each instance (*t*_*s*_, …, *t*_*e*_) of state *n* (see “Cross-syllable likelihoods” section above for a definition of state instances), we inserted the first *t*_*e*_ – *t*_*s*_ frames of *h* into dimensions 5*n*, …,5*n* + 5 of the design matrix, aligning the first frame of *h* to frame *t*_*s*_ in the design matix. Kinematic features were regressed against the design matrix using Ridge regression from scikit-learn and 5-fold cross-validation. We used a range of values from 10^−3^ to 10^3^ for the regularization parameter α and reported the results with greatest accuracy.

### Rearing analysis

To compare the dynamics of rear-associated states across methods, we systematically identified all instances of rearing in our initial open field dataset. During a stereotypical rear, mice briefly stood on their hindlegs and extended their head upwards, leading to a transient increase in height from its modal value of 3cm-5cm to a peak of 7cm-10cm. Rears were typically brief, with mice exiting and then returning to a prone position within a few seconds. We encoded these features using the following criteria. First, rear onsets were defined as increases in height from below 5cm to above 7cm that occurred within the span of a second, with onset formally defined as the first frame where the height exceeded 5cm. Next, rear offsets were defined as decreases in height from above 7cm to below 5cm that occurred within the span of a second, with offset formally defined as the first frame where the height fell below 7cm. Finally, we defined complete rears as onset-offset pairs defining an interval with length between 0.5 and 2 seconds. Height was determined from the distribution of depth values in cropped, aligned and background-segmented videos. Specifically, we used the 98^th^ percentile of the distribution in each frame.

### Accelerometry processing

From the IMU we obtained absolute rotations *r*_*y*_, *r*_*p*_, *r*_*r*_ (yaw, pitch, and roll) and accelerations *a*_*x*_, *a*_*y*_, *a*_*z*_ (dorsal/ventral, posterior/anterior, left/right). To control for subtle variations in implant geometry and chip calibration, we centered the distribution of sensor readings for each variable on each session. We defined total acceleration as the norm of the 3 acceleration components:

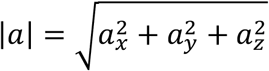

Similarly, we defined total angular velocity as the norm |ω| of rotation derivative:

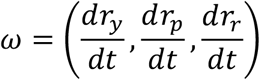

Finally, to calculate jerk, we smoothed the acceleration signal with a 50ms Gaussian kernel, generating a time-series *ã*, and then computed the norm of its derivative:

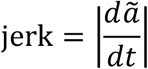

### Aligning dopamine fluctuations to behavior states

For a detailed description of photometry data acquisition and preprocessing, see ref^1^. Briefly, photometry signals were: (1) ΔF/F0-normalized using a 5-second window; (2) adjusted against a reference to remove motion artefacts and other non-ligand-associated fluctuations; (3) z-scored using a 20-second sliding window; (4) temporally aligned to the 30Hz behavioral videos.

Given a set of state onsets (either for a single state or across all states), we computed the onset-aligned dopamine trace by averaging the dopamine signal across onset-centered windows. From the resulting traces, each of which can be denoted as a time-series of dopamine signal values (*d*_−*T*_, …, *d*_*T*_) we defined the total fluctuation size (Fig 5d) and temporal asymmetry (Fig 5e) as

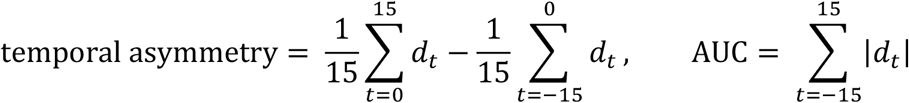

A third metric – the average dopamine during each state (Extended Data Figure 6b) – was defined simply as the mean of the dopamine signal across all frames bearing that state label. For each metric, shuffle distributions were generated by repeating the calculation with a temporally reversed copy of the dopamine times-series.

### Supervised behavior benchmark

Videos and behavioral annotations for the supervised open field behavior benchmark (Fig 4a-c) were obtained from (Bohnslav, 2019)^11^. The dataset contains 20 videos that are each 10-20 minutes long. Each video includes frame-by-frame annotations of five possible behaviors: locomote, rear, face groom, body groom, and defecate. We excluded “defecate” from the analysis because it was extremely rate (< 0.1% of frames).

For pose tracking we used DLC’s SuperAnimal inference API that performs inference on videos without the need to annotate poses in those videos. Specifically, we used SuperAnimal-TopViewMouse that applies DLCRNet-50 as the pose estimation model11. Keypoint detections were obtained using DeepLabCut’s API function deeplabcut.video_inference_superanimal. The API function uses a pretrained model called SuperAnimal-TopViewMouse and performs video adaptation that applies multi-resolution ensemble (i.e., the image height resized to 400, 500, 600 with a fixed aspect ratio) and rapid self-training (model trained on zero shot predictions with confidence above 0.1) for 1000 iterations to counter domain shift and reduce jittering predictions. The code to reproduce this analysis is:

~~~
   videos = [‘path_to_video’]
   superanimal_name = ‘superanimal_topviewmouse’
   scale_list = [400, 500, 600]
   deeplabcut.video_inference_superanimal(videos,
      superanimal_name,
      videotype=“.mp4”,
      video_adapt = True,
      scale_list = scale_list)
~~~

Keypoint coordinates and behavioral annotations for the supervised social behavior benchmark (Fig 4d-f) were obtained from the CalMS21 dataset^12^ (task1). The dataset contains 70 videos of resident-intruder interactions with frame-by-frame annotations of four possible behaviors: attack, investigate, mount, or other. All unsupervised behavior segmentation methods were fit to 2D keypoint data for the resident mouse.

We used four metrics^9^ to compare supervised annotations and unsupervised states from each method. These included normalized mutual information, homogeneity, adjusted rand score, and purity. All metrics besides purity were computed using the python library scikit-learn (i.e., with the function normalized_mutual_info_score, homogeneity_score, adjusted_rand_score). The purity score was defined as in ref^9^.

## MATHEMATICAL MODELING

### Notation

1. χ^−2^(ν, τ^2^) denotes the scaled inverse Chi-squared distribution.
2. ⊗ denotes the Kronecker product.
3. Δ^*N*^ is the *N*-dimensional simplex.
4. *I*_*N*_ is the *N* × *N* identity matrix.
5. 1_*N*×*M*_ is the *N* × *M* matrix of ones.
6. 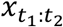 denotes the concatenation 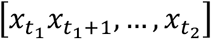 where *t*_1_ < *t*_2_.

### Generative model

Keypoint-MoSeq learns syllables by fitting a switching linear dynamical systems (SLDS) model^13^, which decomposes an animal’s pose trajectory into a sequence of stereotyped dynamical motifs. In general, SLDS models explain time-series observations *y*_1_, …, *y*_*T*_ through a hierarchy of latent states, including continuous states *x*_*t*_ ∈ ℝ^*M*^ that represent the observations *y*_*t*_ in a low-dimensional space, and discrete states *z*_*t*_ ∈ {1, …, *N*} that govern the dynamics of *x*_*t*_ over time. In keypoint-MoSeq, the discrete states correspond to syllables, the continuous states correspond to pose, and the observations are keypoint coordinates. We further adapted SLDS by (1) including a sticky Hierarchical Dirichlet prior (HDP); (2) excplicitly modeling the animal’s location and heading; (3) including a robust (heavy-tailed) observation distribution for keypoints. Below we review SLDS models in general and then describe each of the customizations implemented in keypoint-MoSeq.

#### Switching linear dynamical systems

The discrete states *z*_*t*_ ∈ {1, …, *N*} are assumed to form a Markov chain, meaning

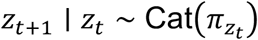

where π_*i*_ ∈ Δ^*N*^ is the probability of transitioning from discrete state *i* to each other state. Conditional on the discrete states *z*_*t*_, the continuous states *x*_*t*_ follow an *L*-order vector autoregressive process with Gaussian noise. This means that the expected value of each *x*_*t*_ is a linear function of the previous *L* states *x*_*t*−*L*:*t*−1_, as shown below,

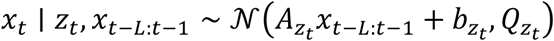

where *A*_*i*_ ∈ ℝ^*M*×*LM*^ is the autoregressive dynamics matrix, *A*_>_ ∈ ℝ^*M*^ is the dynamics bias vector, and *Q*_*i*_ ∈ ℝ^*M*×*M*^ is the dynamics noise matrix for each discrete state *i* = 1, …, *N*. The dynamics parameters (*A*_*i*_, *b*_*i*_, *Q*_*i*_) have a matrix normal inverse Wishart (MNIW) prior,

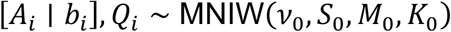

where ν_0_ > *M* − 1 is the degrees of freedom, *S*_0_ ∈ ℝ^*M*×*M*^ is the prior covariance matrix, *M*_0_ ∈ ℝ^*M*×(*LM*+1)^ is the prior mean dynamics matrix, and *K*_0_ ∈ ℝ^(*LM*+1)×(*LM*+1)^ is the prior scale matrix. Finally, in the standard formulation of SLDS (which we modify for keypoint data, as described below), each observation *y*_*t*_ ∈ ℝ^*D*^ is a linear function of *x*_*t*_ plus noise:

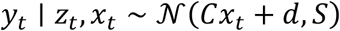

Here we assume that the observation parameters *C, d* and *S* do not depend on *z*_*t*_.

#### Sticky hierarchical Dirichlet prior

A key feature of depth Moseq^7^ is the use of a sticky HDP prior^14^ for the transition matrix. In general, HDP priors allow the number of distinct states in a hidden Markov model to be inferred directly from the data. The “sticky” variant of the HDP prior includes an additional hyper-parameter κ that tunes the frequency of self-transitions in the discrete state sequence *z*_*t*_, and thus the distribution of syllable durations. As in depth MoSeq, we implement a sticky-HDP prior using the weak limit approximation^14^, as shown below:

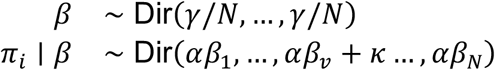

where κ is being added in the *i*th position. Here β ∈ *Δ*^*N*^ is a global vector of augmented syllable transition probabilities, and the hyperparameters γ, α, κ control the sparsity of states, the weight of the sparsity prior, and the bias toward self-transitions respectively.

#### SLDS for postural dynamics

Keypoint coordinates reflect not only the pose of an animal, but also its location and heading. To disambiguate these factors, we define a canonical, egocentric reference frame in which the postural dynamics are modeled. The canonically aligned poses are then transformed into global coordinates using explicit centroid and heading variables that are learned by the model.

Concretely, let *Y*_*t*_ ∈ ℝ^*K*×*D*^ represent the coordinates of *K* keypoints at time *t*, where *D* ∈ {2,*3*}. We define latent variables *v*_*t*_ ∈ ℝ^*D*^ and *h*_*t*_ ∈ [0,2π] to represent the animal’s centroid and heading angle. We assume that each heading angle *h*_*t*_ has an independent, uniform prior and that the centroid is autocorrelated as follows:

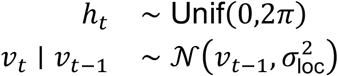

At each time point *t*, the pose *Y*_*t*_ is generated via rotation and translation of a centered and oriented pose 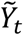 that depends on the current continuous latent state *x*_*t*_:

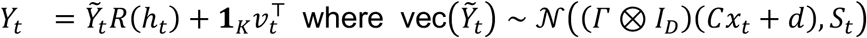

where *R(h*_*t*_) is a matrix that rotates by angle *h*_*t*_ in the xy-plane, and Γ ∈ *R*^*K*×(*K*−1)^ is defined by the truncated singular value decomposition Γ Δ Γ^*T*^ = *I*_*K*_ – **1**_*K*×*K*_*/K*. Note that Γ encodes a linear transformation that isometrically maps ℝ^(*K*−1)×*D*^ to the set of all centered keypoint arrangements in ℝ^*K*×*D*^, and thus ensures that 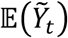 is always centered^15^. The parameters *C* ∈ ℝ^(*K*−1)*D*×*M*^, and *d* ∈ ℝ^(*K*−1)*D*^ are initialized using principal components analysis (PCA) applied to the transformed keypoint coordinates 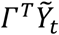. In principle *C* and *d* can be adjusted further during model fitting, and we describe the corresponding Gibbs updates in the inference section below. In practice, however, we keep *C* and *d* fixed to their initial values when fitting keypoint-MoSeq.

#### Robust observations

To account for occasional large errors during keypoint tracking, we use the heavy-tailed Student’s *t*-distribution, which corresponds to a normal distribution whose variance is itself a random variable. Here, we instantiate the random variances explicitly as a product of two parameters: a baseline variance *σ*_*k*_ for each keypoint and a time-varying scale *s*_*t,k*_. We assume:

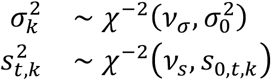

where ν_σ_ > 0 and ν_*s*_ > 0 are degrees of freedom, 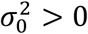 is a baseline scaling parameter, and *s*_0,*t,k*_ > 0 is a local scaling parameter, which encodes a prior on the scale of error for each keypoint on each frame. Where possible, we calculated the local scaling parameters as a function of the neural network confidences for each keypoint. The function was calibrated using the empirical relationship between confidence values and error sizes. The overall noise covariance *S*_*t*_ is generated from σ_*k*_ and *s*_*t,k*_ as follows:

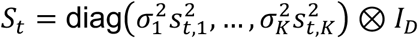

#### Related work

Keypoint-MoSeq extends the model used in depth MoSeq^7^, where a low-dimensional pose trajectory *x*_*t*_ (derived from egocentrically aligned depth videos) is used to fit an autoregressive hidden Markov model with a transition matrix π, autoregressive parameters *A*_*i*_, *b*_*i*_, *Q*_*i*_ and discrete states *z*_*t*_ like those described here. Indeed, conditional on *x*_*t*_, the models for keypoin-MoSeq and depth MoSeq are identical. The main differences are that keypoint-MoSeq treats *x*_*t*_ as a latent variable (i.e. updates it during fitting), includes explicit centroid and heading variables, and uses a robust noise model.

Disambiguating pose from position and heading is a common task in unsupervised behavior algorithms, and researchers have adopted a variety of approaches. VAME^9^, for example, isolates pose by centering and aligning data ahead of time, whereas B-SOiD^8^ transforms the keypoint data into a vector of relative distances and angles. The statistical pose model GIMBAL^5^, on the other hand, introduces latent heading and centroid variables that are inferred simultaneously with the rest of the model. Keypoint-MoSeq adopts this latter approach, which is able to remove spurious correlations between egocentric features that can arise from errors in keypoint localization.

### Inference algorithm

Our full model contains latent variables *v, h, x, z, s* and parameters *A, b, Q, C, d*, σ, β, π. We fit each of these variables – with the exception of *C* and *d* – using Gibbs sampling, in which each variable is iteratively resampled from its posterior distribution conditional on the current values of all the other variables. The posterior distributions *P*(π, β ∣ *z*) and *P(A, b, Q* ∣ *z, x*) are unchanged from the original MoSeq paper and will not be be reproduced here (see ref^7^, pages 42-44, and note the changes of notation *Q* → Σ, *z* → *x*, and *x* → *y*). *h* are described below.

#### Resampling P(C, d ∣ s, σ, x, v, h, Y)

Let 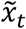 represent *x*_*t*_ with a 1 appended and define

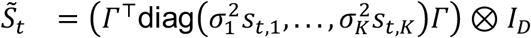

The posterior update is (*C, d*) *∼𝒩* (vec(*C, d*) ∣ μ_*n*_, *Σ*_*n*_) where

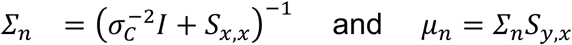

with

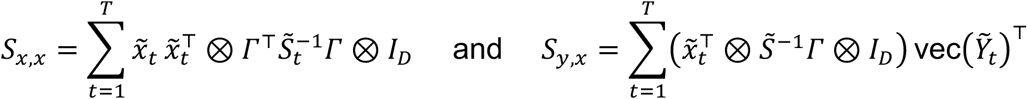

#### Resampling P(s ∣ C, d, σ, x, v, h, Y)

Each *s*_*t,k*_ is conditionally independent with posterior

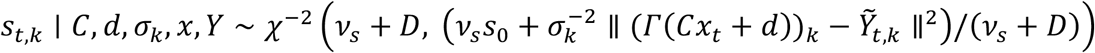

#### Resampling P(σ ∣ C, d, s, x, v, h, Y)

Each σ_*k*_ is conditionally independent with posterior

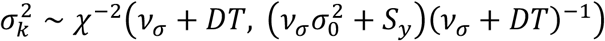

where 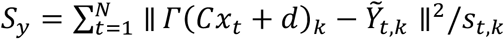

#### Resampling P(v∣ C, d, σ, s, x, h, Y)

Since the translations *v*_1_, …, *v*_*T*_ form a linear dynamical system, they can be updated by Kalman sampling. The observation potentials have the form 𝒩(*v*_*t*_ ∣ μ, γ^2^*I*_*D*_) where

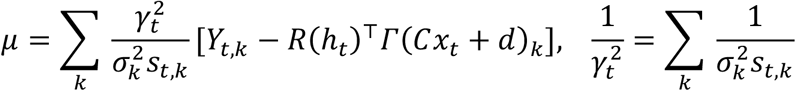

#### Resampling P(h ∣ C, d, σ, s, x, v, Y)

The posterior of *h*_*t*_ is the von-Mises distribution vM(θ, κ) where κ and θ ∈ [0,2π] are the unique parameters satisfying [κcos(θ), κsin(θ)] = [*S*_1,1_ + *S*_2,2_, *S*_1,2_ – *S*_2,1_] for

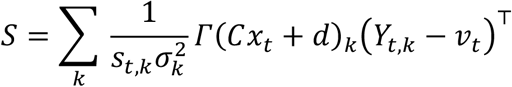

#### Resampling P(x ∣ C, d, σ, s, v, h, Y)

To resample *x*, we first express its temporal dependencies as a first-order autoregressive process, and then apply Kalman sampling. The change of variables is

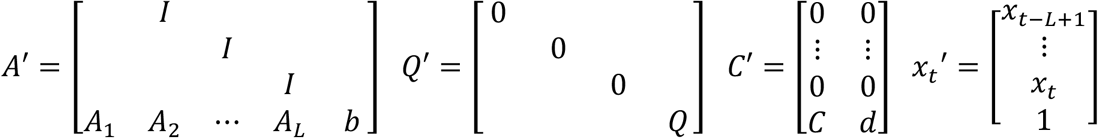

Kalman sampling can then be applied to the sample the conditional distribution,

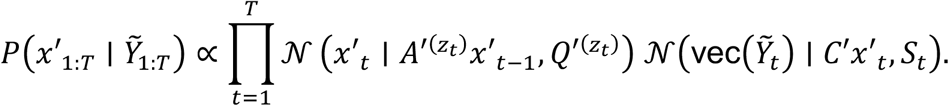

(Assume *x*′ is left-padded with zeros for negative time indices.)

### Hyper-parameters

We used the following hyper-parameter values throughout the paper.

#### Transition matrix

*N* = 100

*γ* = 1000

*α* = 100

*κ* fit to each dataset

#### Autoregressive process

*M* set using PCA explained variance curve

*L* = *3*

*ν*_0_ = *M* + 2

*S*_0_ = 0.01*I*_*M*_

*M*_0_ = [0_*M*×(*L*−1)_ *I*_*M*_ 1_*M*×1_]

*K*_0_ = 10*I*_*M(L*+1)_

#### Observation process

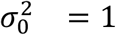

*ν*_*σ*_ = 10^5^

*ν*_*s*_ = 5

*S*_0,*t,k*_ set based on neural network confidence

#### Centroid autocorrelation

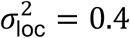

### Derivation of Gibbs updates

#### Derivation of C, d updates

To simply notation, define

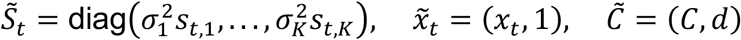

The likelihood of the centered and aligned keypoint locations 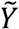 can be expanded as follows.

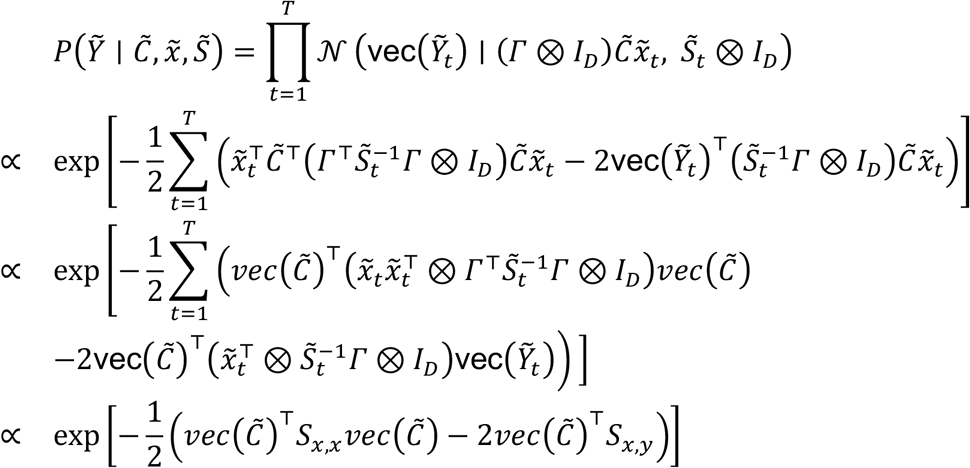

where

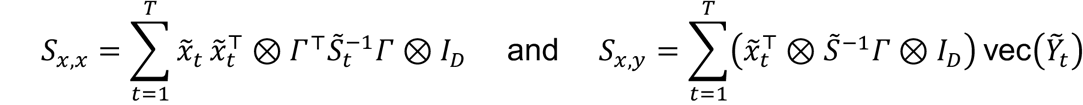

Multiplying by the prior 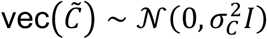 yields

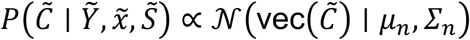

where

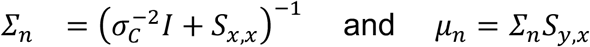

#### Derivation of σ_k_, s_t,k_ updates

For each time *t* and keypoint *k*, let 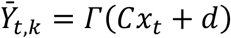. The likelihood of the centered and aligned keypoint location 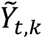 is

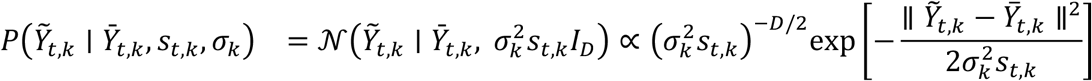

We can then calculate posteriors *P(s*_*t,k*_ ∣ σ_*k*_) and *P*(σ_*k*_ ∣ *s*_*t,k*_) as follows.

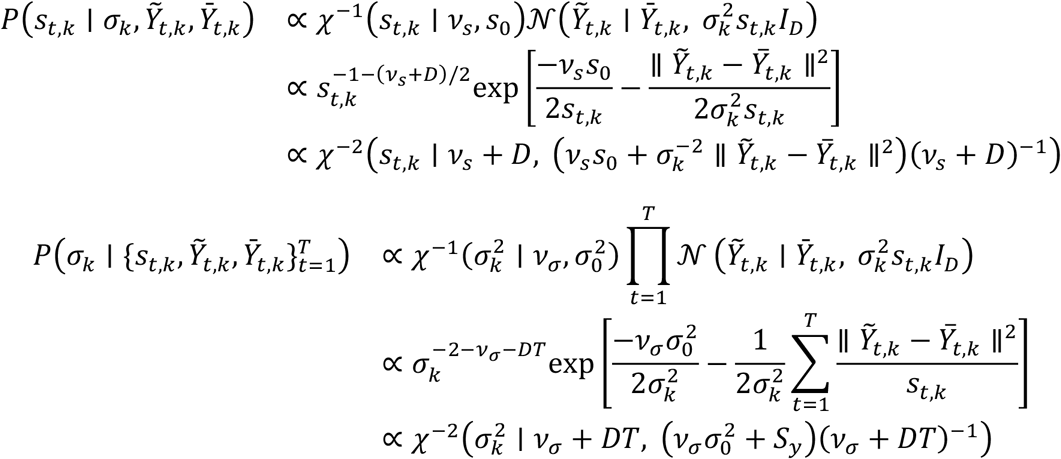

where 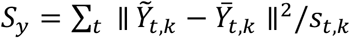

#### Derivation of v_t_ update

We assume an improper uniform prior on *v*_*t*_, hence

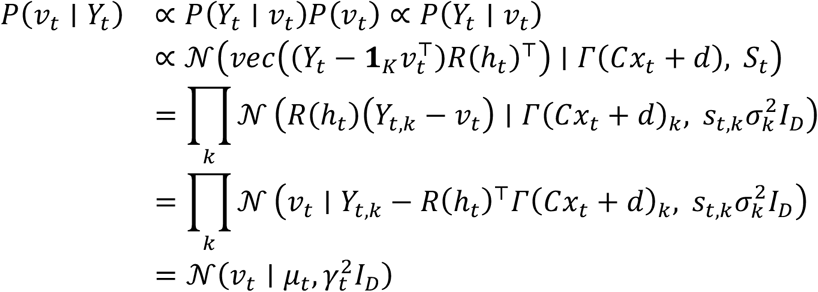

where

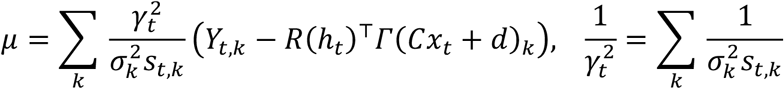

#### Derivation of h_t_ update

We assume a proper uniform prior on *h*_*t*_, hence

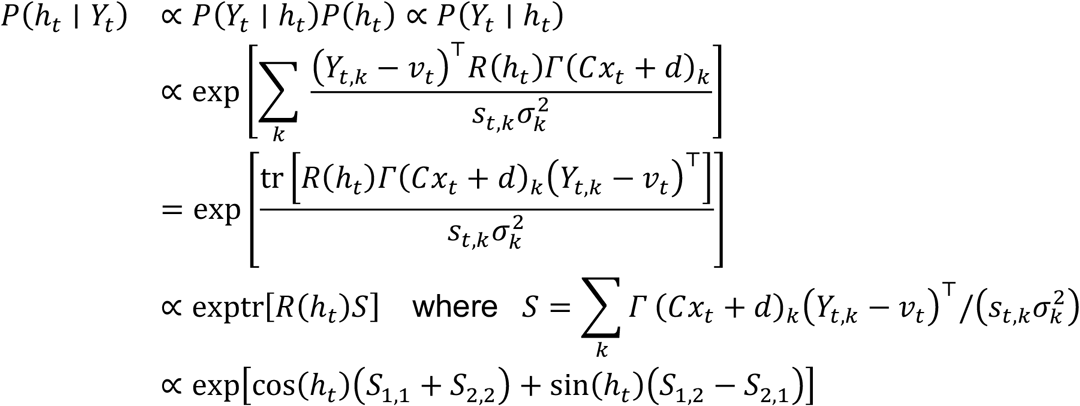

Let [κco*s*(θ), κsin(θ)] represent [*S*_1,1_ + *S*_2,2_, *S*_1,2_ – *S*_2,1_] in polar coordinates. Then

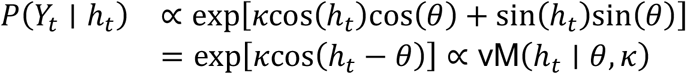

## Extended Data

**Extended Data Figure 1:**
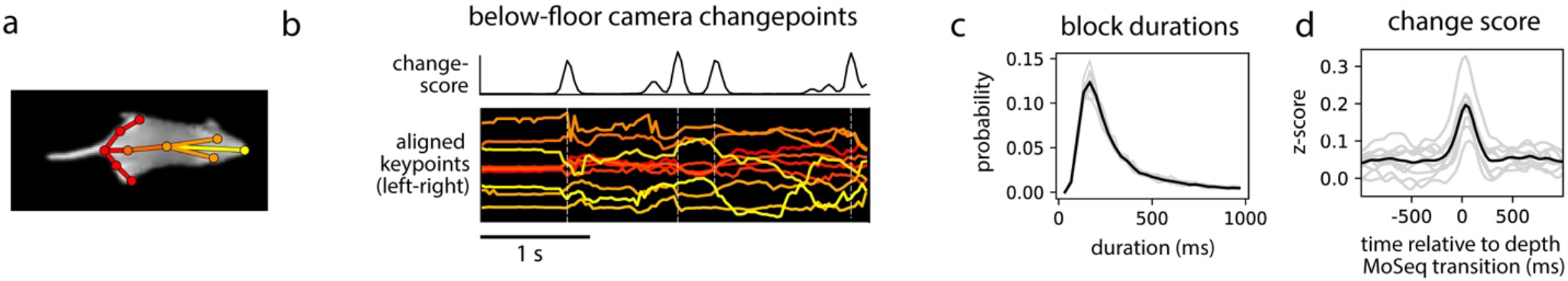
Mouse behavior exhibits sub-second syllable structure when keypoints are tracked from below. **a)** 2D keypoints tracked using infrared video from a camera viewing the mouse through a transparent floor. **b)** Egocentrically aligned keypoint trajectories (bottom) and change scores derived from those keypoints (top, see Methods). Vertical dashed lines represent changepoints (peaks in the change score). **c)** Distribution of inter-changepoint intervals. **d)** Keypoint change score aligned to syllable transitions from depth MoSeq. Results in (c) and (d) are shown for the full dataset (black lines) and for each recording session (gray lines).

**Extended Data Figure 2:**
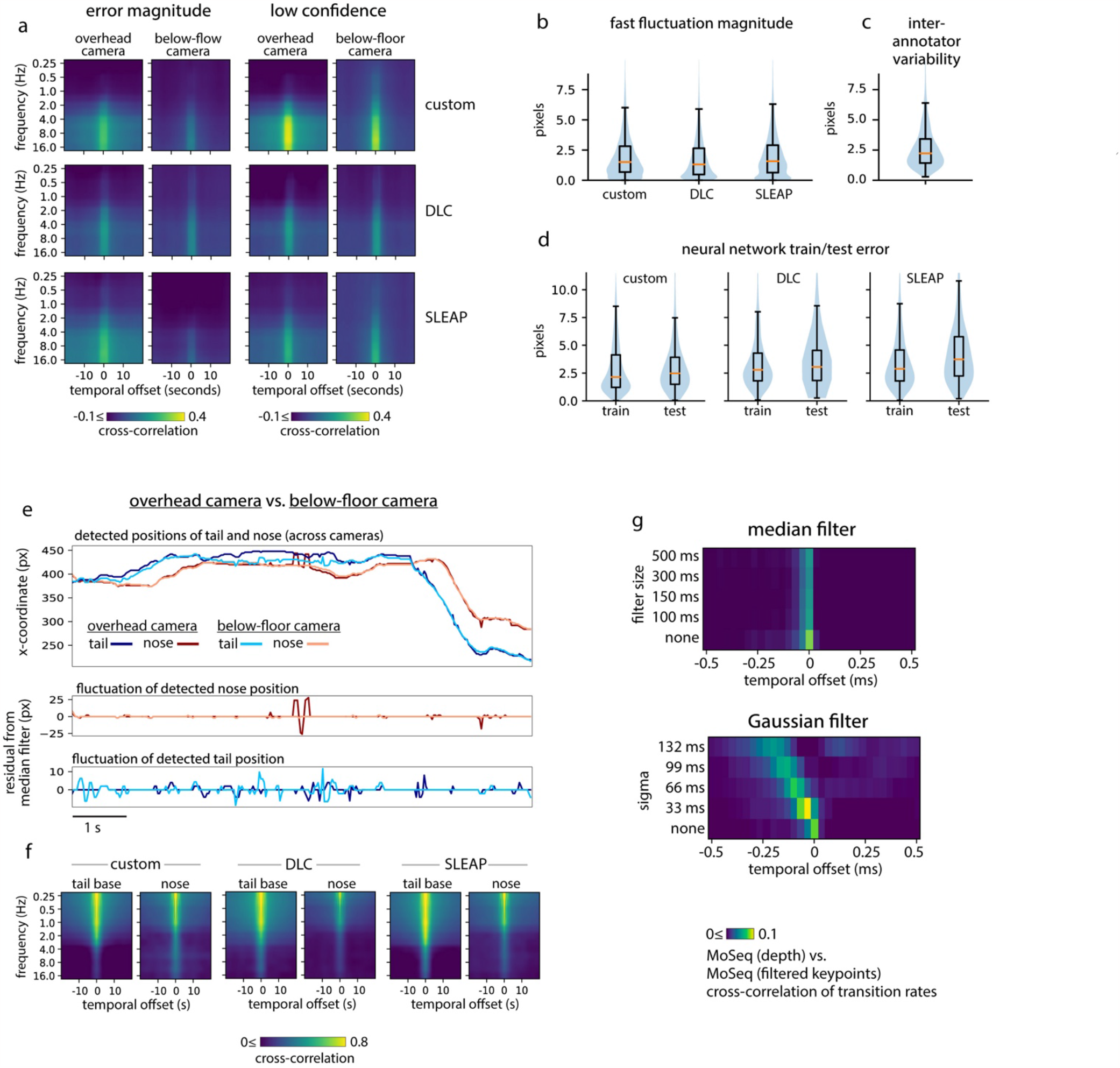
Markerless pose tracking exhibits fast fluctuations are that are independent of behavior yet affect MoSeq output. **a) Noise-driven fast fluctuations are pervasive across camera angles and tracking methods**. Cross-correlation between the spectral content of keypoint fluctuations and either error magnitude (left) or a measure of low-confidence keypoint detections (right) (see Methods). **b-d) Tracking noise reflects ambiguity in keypoint locations. b)** Magnitude of fast fluctuations in keypoint position for three different tracking methods, calculated as the per-frame distance from the measured trajectory of a keypoint to a smoothened version of the same trajectory, where smoothing was performed using a gaussian kernel with width 100ms. **c)** Inter-annotator variability, shown as the distribution of distances between different annotations of the same keypoint. **d)** Train- and test-error distributions for each keypoint tracking method. **e) Fast fluctuations are weakly correlated between camera angles. Top:** position of the nose and tail-base over a 10-second interval, shown for both the overhead and below-floor cameras. **Bottom:** fast fluctuations in each coordinate, obtained as residuals after median filtering. **f)** Cross-correlation between spectrograms obtained from two different camera angles for either the tail base or the nose, shown for each tracking method. **g) Filtering keypoint trajectories does not improve MoSeq output**. Cross-correlation of transitions rates, comparing MoSeq (depth) and MoSeq applied to keypoints with various levels of smoothing using either a Gaussian or median filter.

**Extended Data Figure 3:**
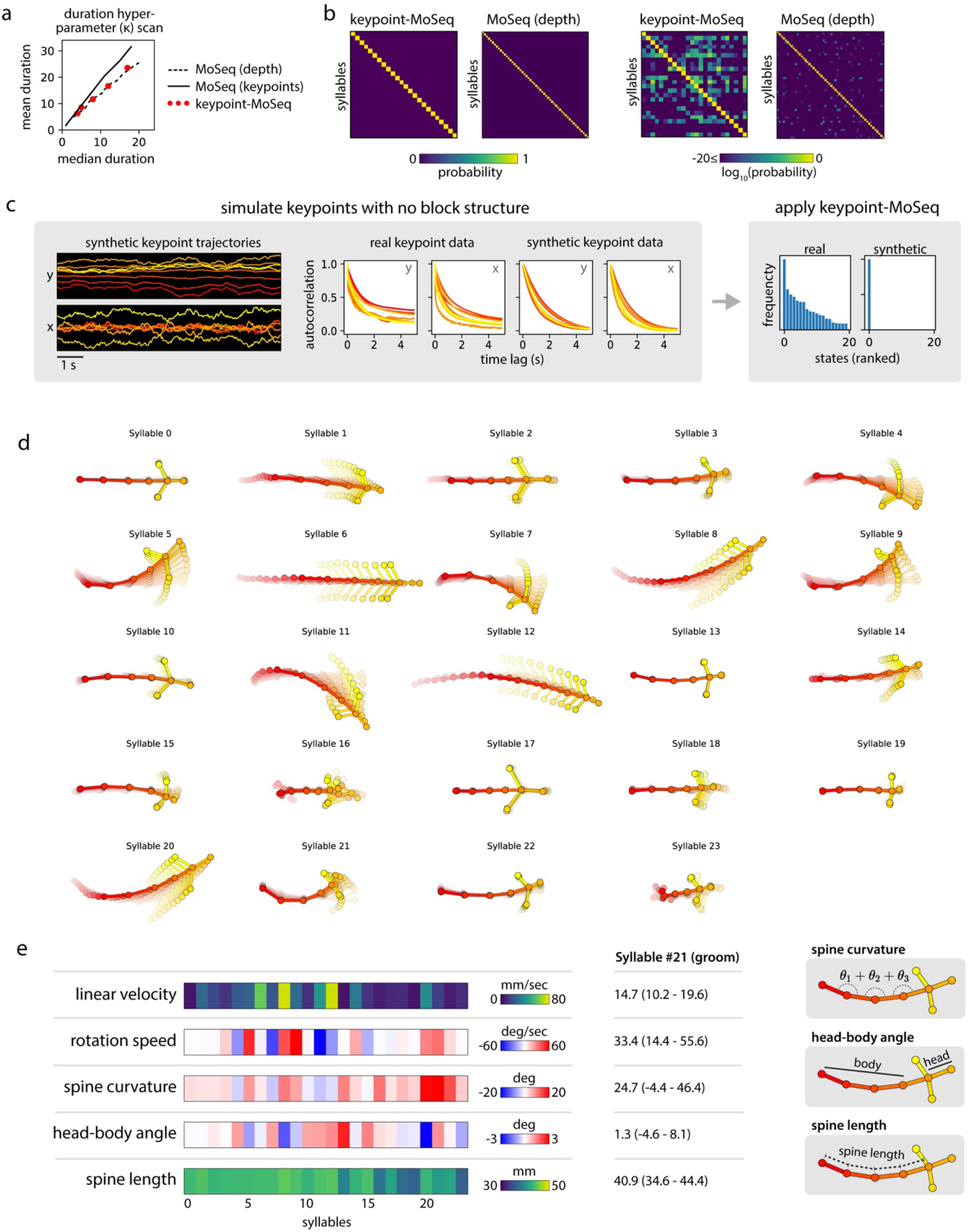
Keypoint-MoSeq partitions behavior into distinct, well-defined syllables. **a) Keypoint-MoSeq and depth MoSeq yield similar duration distributions**. Relationship between mean and median syllable duration as the temporal stickiness hyper-parameter *κ* is varied, shown for keypoint-MoSeq (red dots), as well as original MoSeq applied to depth (dashed line) or keypoints (solid line). **b) Keypoint-MoSeq syllables represent distinguishable pose trajectories**. Syllable cross-likelihoods, defined as the probability, on average, that time-intervals assigned to one syllable (column) could have arisen from another syllable (row). Cross-likelihoods were calculated for keypoint-MoSeq and for depth MoSeq. The results for both methods are plotted twice, using either an absolute scale (left) or a log scale (right). Note that the off-diagonal cross-likelihoods apparent for keypoint-MoSeq on the log scale are practically negligible; we show them here to emphasize that MoSeq models have higher uncertainty when fed lower dimensional data like keypoints compared to depth data. **c) Keypoint-MoSeq fails to distinguish syllables when input data lacks changepoints**. Modeling results for synthetic keypoint data with a similar statistical structure as the real data but lacking in changepoints (see Methods). **Left:** example of synthetic keypoint trajectories. **Middle:** autocorrelation of keypoint coordinates for real vs. synthetic data, showing similar dynamics at short timescales. **Right:** distribution of syllable frequencies for keypoint-MoSeq models trained on real vs. synthetic data. **d-e) Syllable-associated kinematics. d)** Average pose trajectories for syllables identified by keypoint-MoSeq. Each trajectory includes ten evenly timed poses from 165ms before to 500ms after syllable onset. **e)** Kinematic and morphological parameters for each syllable.

**Extended Data Figure 4:**
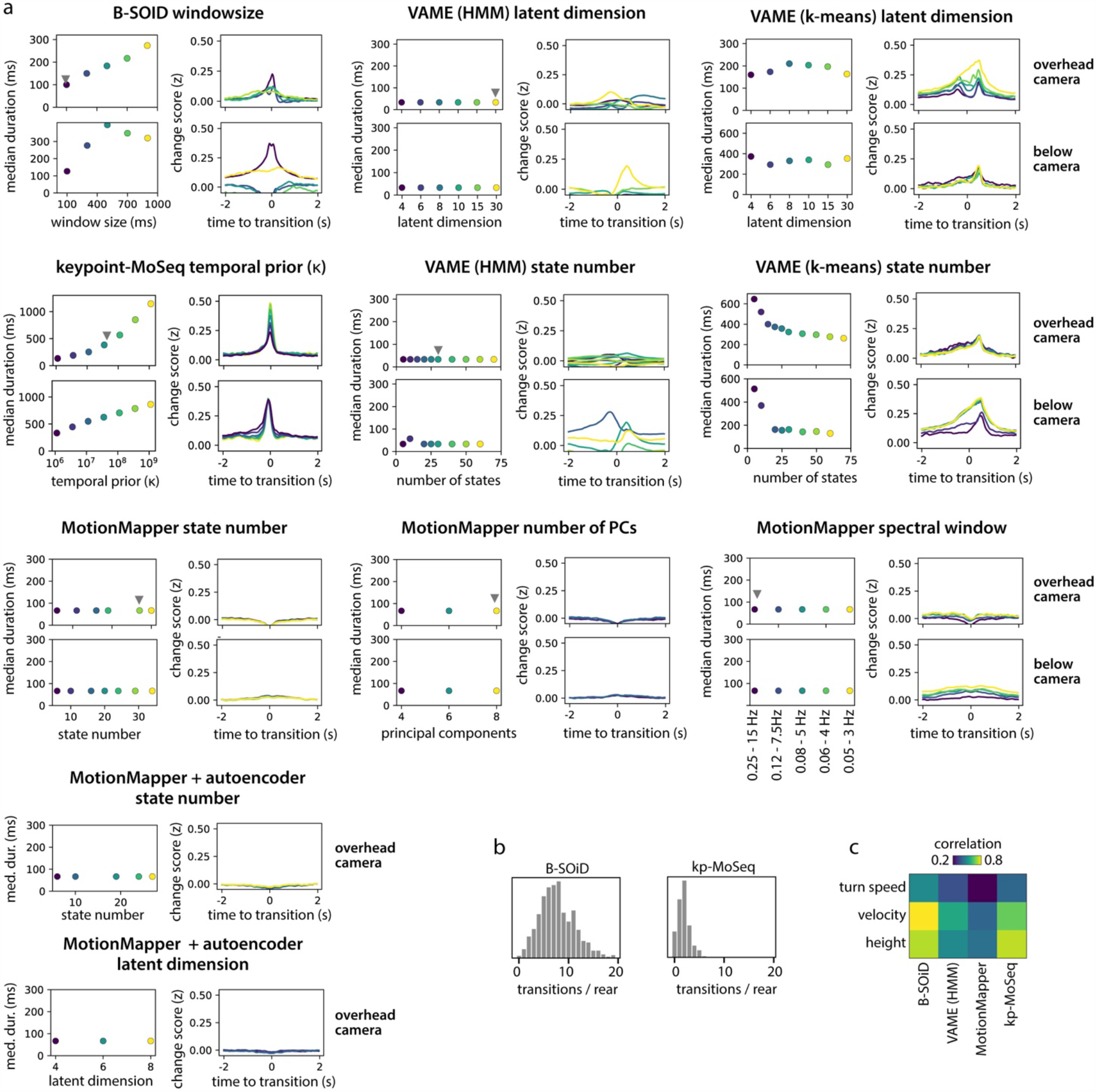
Method-to-method differences in sensitivity to behavioral changepoints are robust to parameter settings. **a)** Output of unsupervised behavior segmentation algorithms across a range of parameter settings, applied to 2D keypoint data from two different camera angles. The median state duration (left) and the average (z-scored) keypoint change score aligned to state transitions (right) are shown for each method and parameter value. Gray pointers indicate default parameter values used for subsequent analysis. **b)** Distributions showing the number of transitions that occur during each rear. **c)** Accuracy of kinematic decoding models that were fit to state sequences from each method.

**Extended Data Figure 5:**
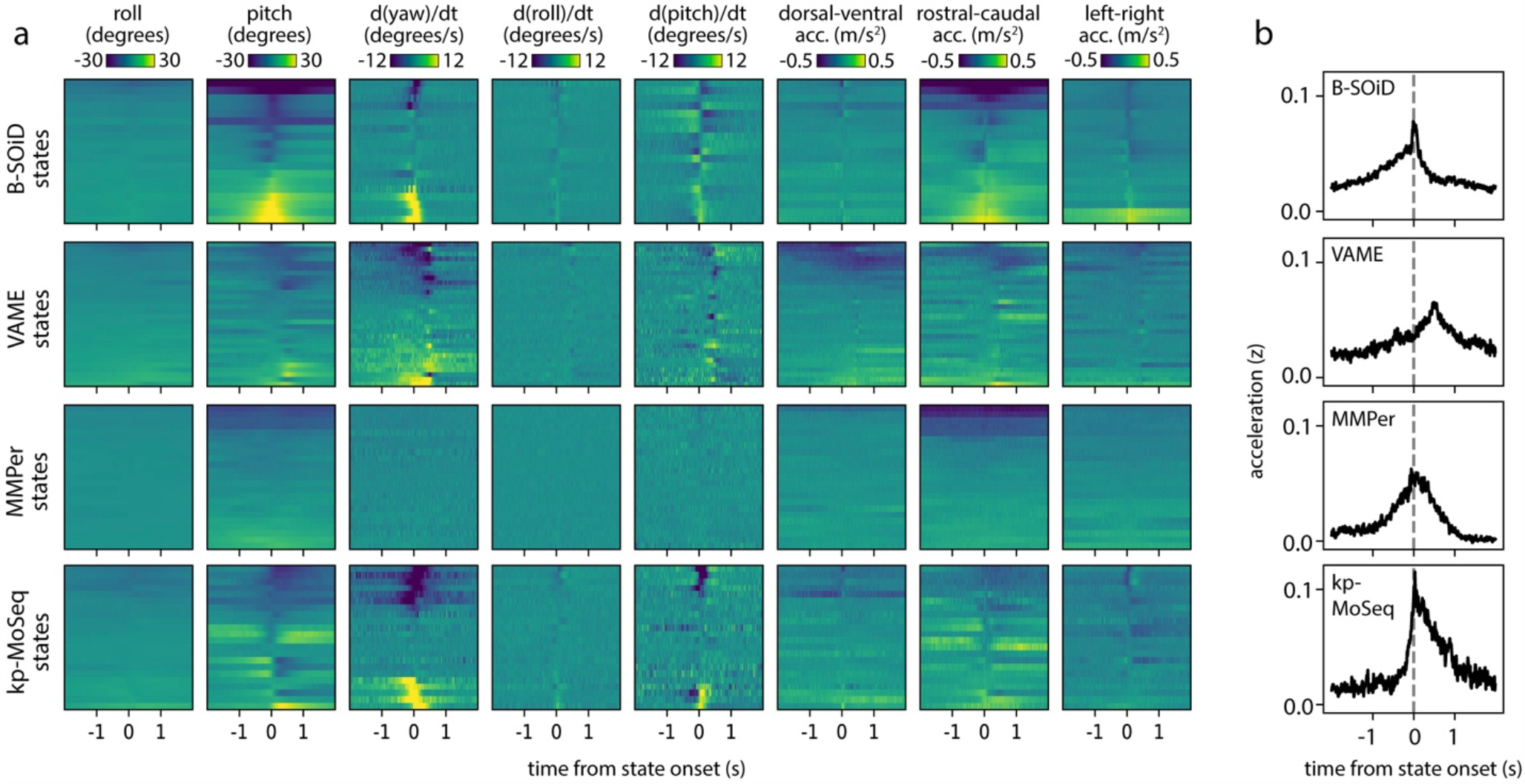
Accelerometry reveals kinematic transitions at the onsets of keypoint-MoSeq states. **a)** IMU signals aligned to state onsets from several behavior segmentation methods. Each row corresponds to a behavior state and shows the average across all onset times for that state. **b)** As (a) for acceleration but showing the median across all states.

**Extended Data Figure 6:**
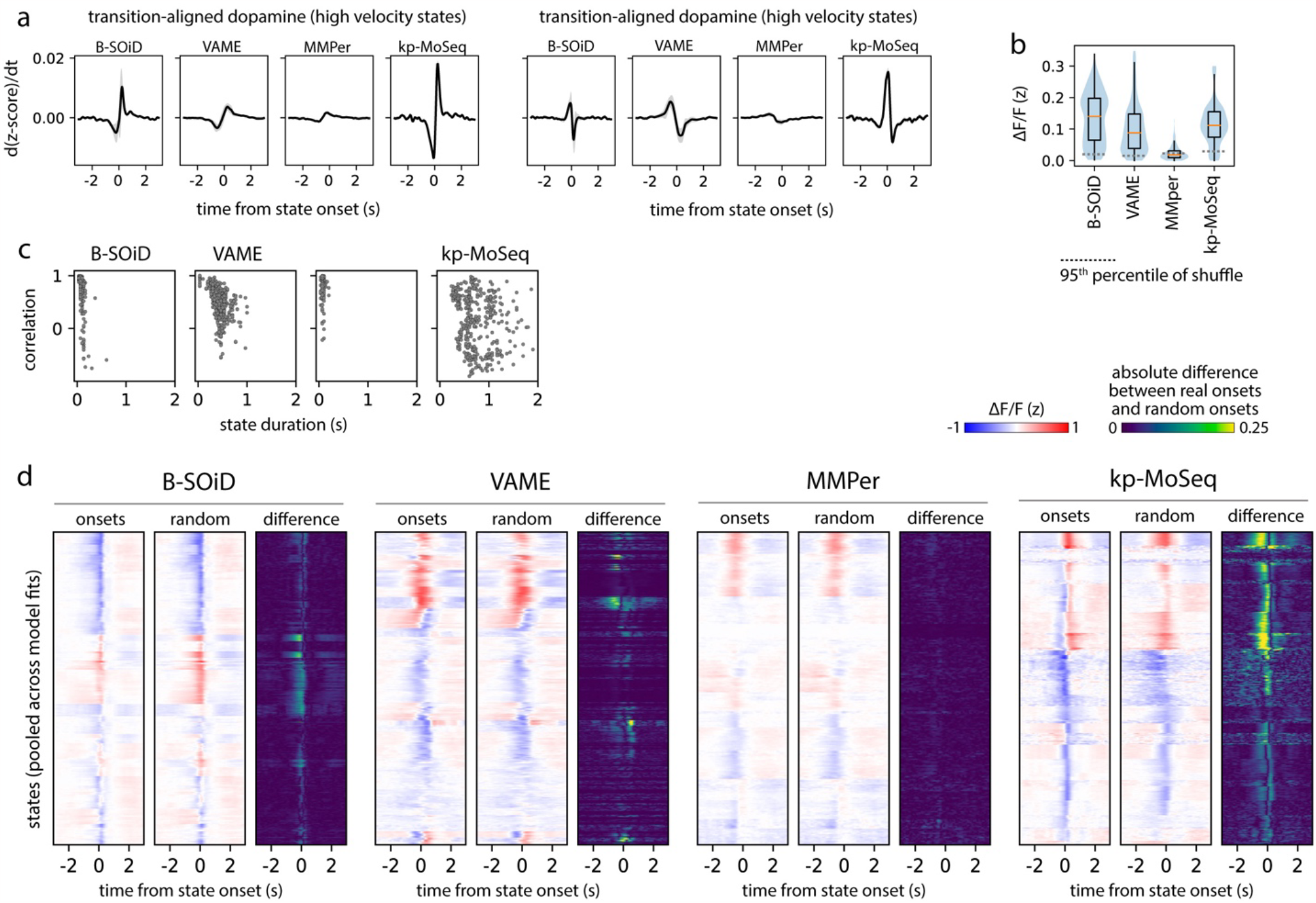
Striatal dopamine fluctuations are enriched at keypoint-MoSeq syllable onsets. **a) Keypoint-MoSeq best captures dopamine fluctuations for both high- and low-velocity behaviors**. Derivative of the dopamine signal aligned to the onsets of high velocity or low velocity behavior states. States from each method were classified evenly as high or low velocity based on the mean centroid velocity during their respective frames. **b)** Distributions capturing the average of the dopamine signal across states from each method. **c-d) Keypoint-MoSeq syllable onsets are meaningful landmarks for neural data analysis. c)** Relationship between state durations and correlations from Fig 5f, showing that the impact of randomization is not a simple function of state duration. **d)** Average dopamine fluctuations aligned to state onsets (left) or aligned to random frames throughout the execution of each state (middle), as well as the absolute difference between the two alignment approaches (right), shown for each unsupervised behavior segmentation approach.

**Extended Data Figure 7:**
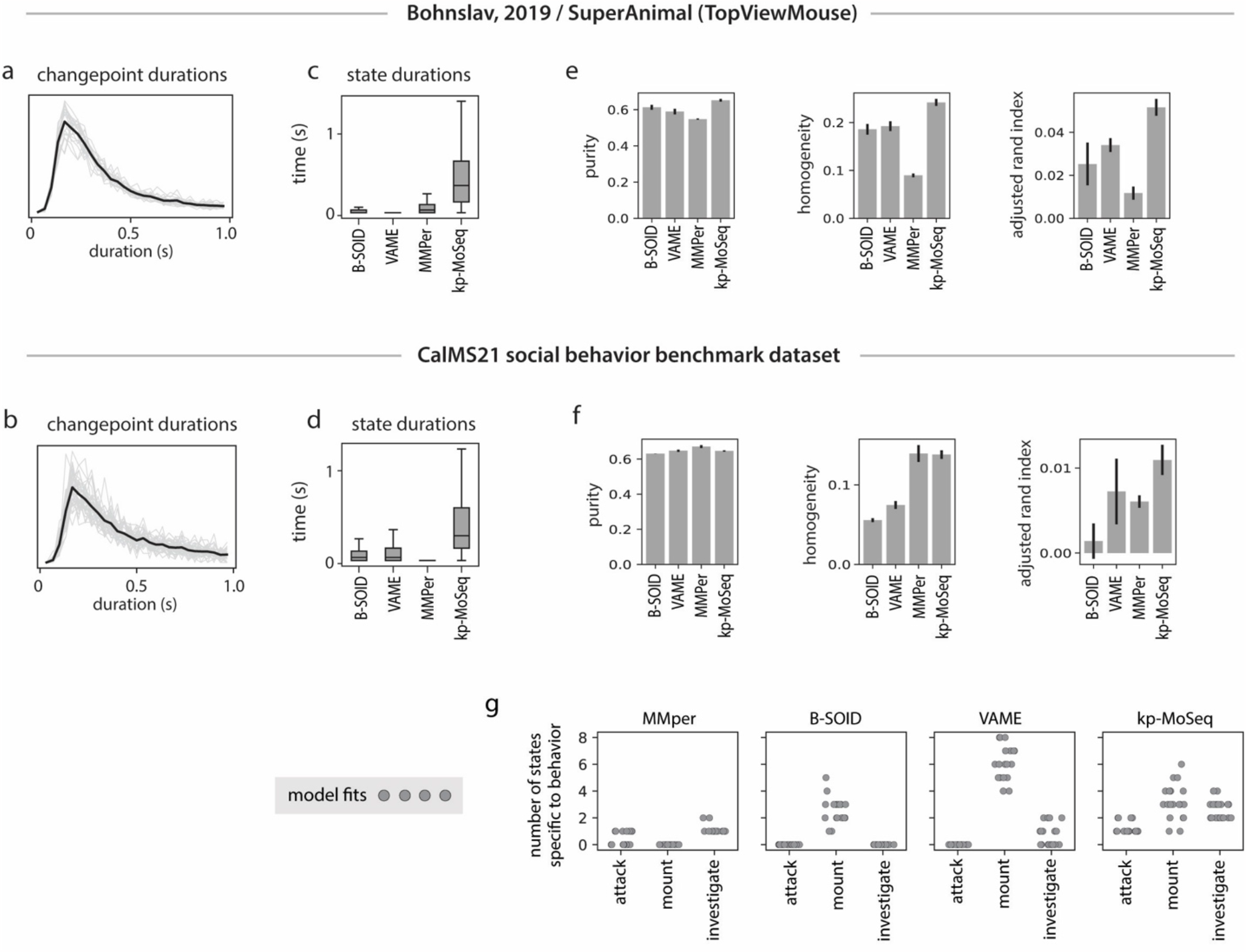
Supervised behavior benchmark. **a-d) Keypoint-MoSeq captures sub-second syllable structure in two benchmark datasets. a**,**b)** Distribution of inter-changepoint intervals for the open field dataset (Bohnslav, 2019) (a) and CalMS21 social behavior benchmark (b), shown respectively for the full datasets (black lines) and for each recording session (gray lines). **c**,**d)** Distribution of state durations from each behavior segmentation method. **e-g) Keypoint-MoSeq matches or outperforms other methods when quantifying the agreement between human-annotations and unsupervised behavior labels. e**,**f)** Three different similarity measures applied to the output of each unsupervised behavior analysis method (see Methods). **g)** Number of unsupervised states specific to each human-annotated behavior in the CalMS21 dataset, shown for 20 independent fits of each unsupervised method. A state was defined as specific if > 50% of frames bore the annotation.

**Extended Data Figure 8:**
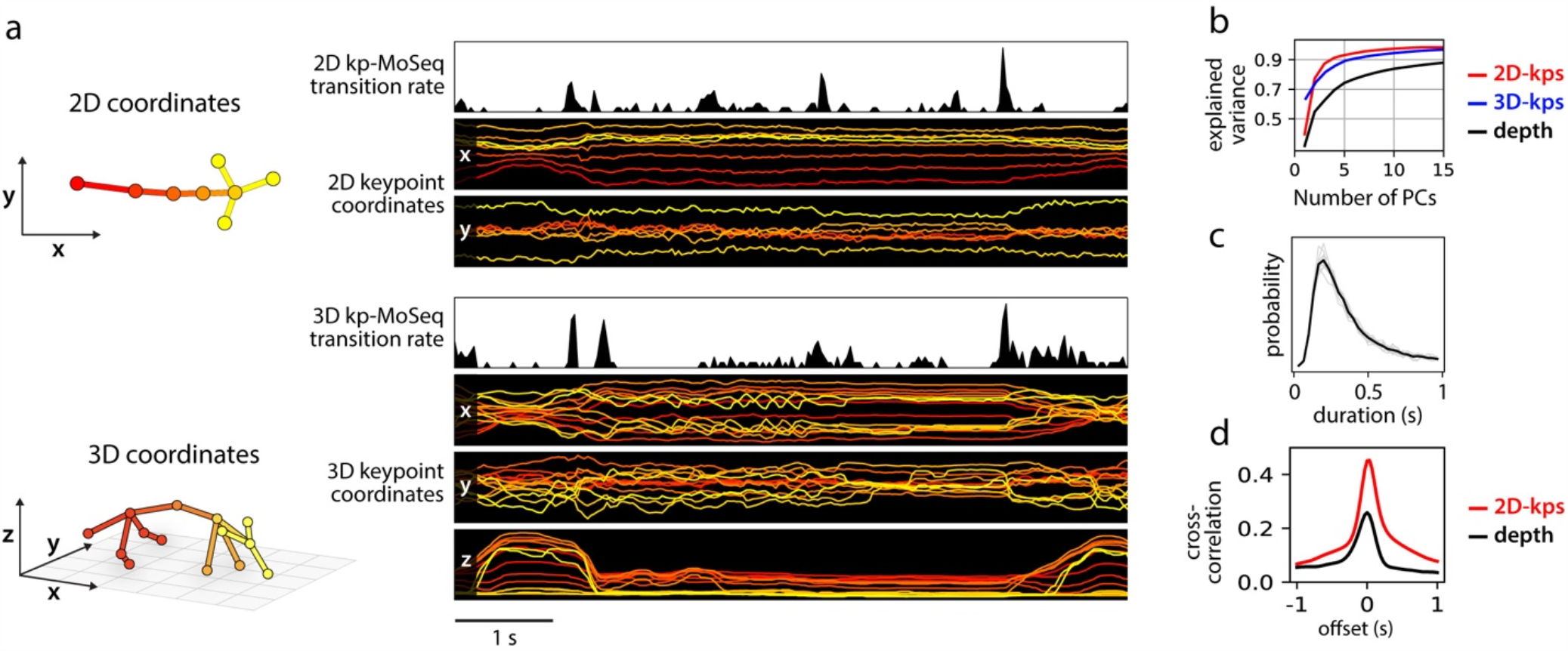
3D and 2D keypoints provide qualitatively distinct pose representations yet share sub-second temporal structure. **a) 3D keypoints have smoother trajectories and exhibit oscillatory gate dynamics. Left:** Keypoints tracked in 2D (top) or 3D (bottom) and corresponding egocentric coordinate axes. Right: example keypoint trajectories and transition rates from keypoint-MoSeq. Transition rate is defined as the posterior probability of a transition occurring on each frame. **b) 2D keypoints, 3D keypoints and depth data provide increasingly high-dimensional pose representations**. Cumulative fraction of explained variance for increasing number of principal components (PCs). PCs were fit to egocentrically aligned 2D keypoints, egocentrically aligned 3D keypoints, or depth videos respectively. **c-d) 3D keypoints capture sub-second syllable structure. c)** Distribution of inter-changepoint intervals in the 3D keypoint dataset, shown. **d)** Cross-correlation between the 3D keypoint change score and change scores derived from 2D keypoints and depth respectively.

**Extended Data Figure 9:**
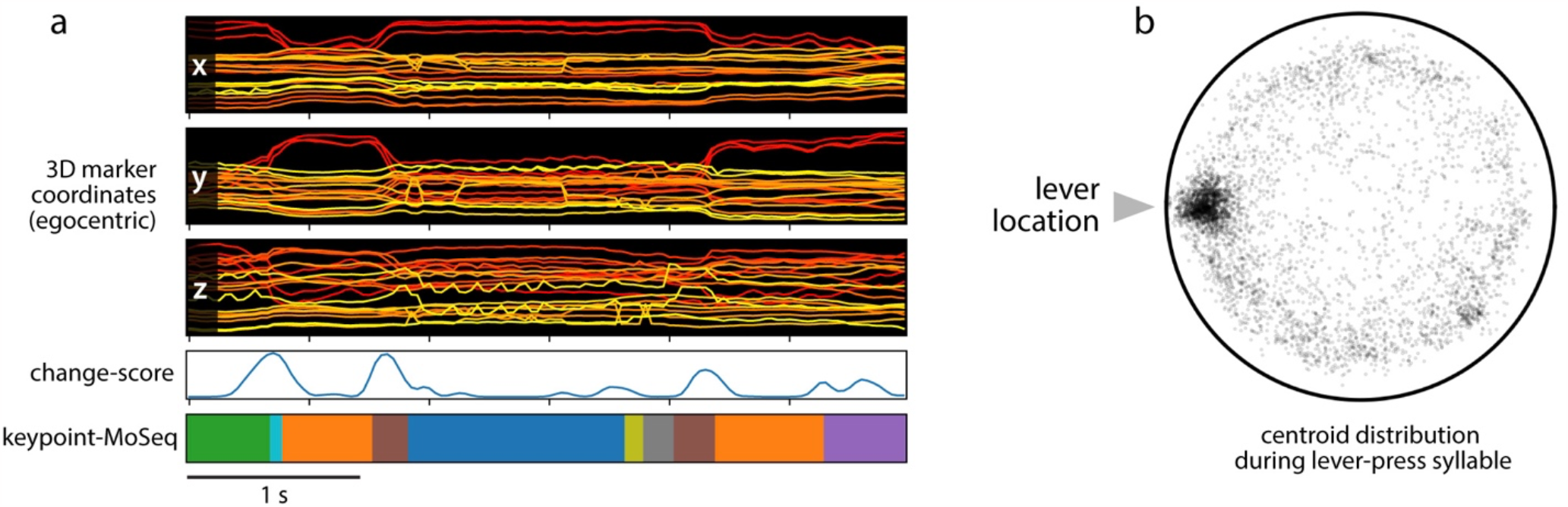
Keypoint-MoSeq analysis of rat motion capture data. **a) Top:** 3D marker positions in egocentric coordinates. **Middle:** change score derived from the marker trajectories. **Bottom:** keypoint-MoSeq syllables. **b)** Random sample of centroid locations during execution of the “lever-press” syllable shown in Fig 6o.

